# Associations between the *Plasmodium falciparum* genome and sickle haemoglobin identified in mild malaria cases from Ghana

**DOI:** 10.1101/2023.09.14.557461

**Authors:** William L. Hamilton, Annie J. Forster, Yaw Aniweh, Victor Asoala, Eleanor Drury, Kirk A. Rockett, Gordon A. Awandare, Dominic P. Kwiatkowski, Gavin Band, Lucas N. Amenga-Etego

**Affiliations:** Wellcome Sanger Institute, Wellcome Trust Genome Campus, Hinxton, CB10 1RQ, United Kingdom; University of Cambridge, Department of Medicine, Cambridge Biomedical Campus, Hills Road, Cambridge CB2 0QQ, United Kingdom; Centre for Human Genetics, Roosevelt Dr, Oxford OX3 7BN, United Kingdom; West African Centre for Cell Biology of Infectious Pathogens (WACCBIP), University of Ghana, Legon, Accra, Ghana; Navrongo Health Research Centre (NHRC), Ghana Health Service, Navrongo, Upper East Region, Ghana

**Keywords:** Malaria, *P. falciparum*, Sickle haemoglobin, HbS, Sickle-associated loci, Genome-wide association, GWAS, host-parasite interaction

## Abstract

**Background:** Heterozygosity for sickle haemoglobin (HbS) confers protection against severe malaria caused by the parasite *Plasmodium falciparum.* Recent work has suggested that this protective effect can depend on the parasite genotype: *P. falciparum* sickle-associated (*Pfsa*) variants were found disproportionately in individuals with severe malaria carrying HbS alleles in The Gambia and Kenya. Interactions between the *P. falciparum* genome and HbS have not previously been investigated in mild malaria cases or in Ghana.

**Methods:** We performed a genome-wide association analysis of *P. falciparum* against human β-globin genotypes in a sample of 1,368 people with mild malaria in northern Ghana.

**Results:** We replicated the previously identified associations with HbS at two parasite loci (*Pfsa1* and *Pfsa3*). *Pfsa2* was absent from this population. A candidate newly identified locus within the serine/ threonine kinase *FIKK4.2*, which we putatively term *Pfsa4*, was also associated with HbS; this finding replicated in a published sample from Mali. The *Pfsa1-4* mutations vary widely in frequencies across Africa, are absent or very low frequency in Asia, and are highly correlated with each-other across multiple populations. We found no strong associations with haemoglobin C.

**Conclusions:** This study replicates previously reported sickle-associated loci in the *P. falciparum* genome and has produced new evidence of a potential association with sickle haemoglobin at a fourth parasite locus. Further research is needed to validate the tentative fourth locus. These findings add new complexity to the emerging picture of association between human and co-evolving malaria parasite genomes, suggesting new avenues for functional exploration.

## Background

The World Health Organization (WHO) estimates that there were 282 million malaria cases and 610,000 deaths in 2024 [1]. 265 million cases (94%) and 579,000 deaths (95%) occurred in the WHO African Region. Just over 75% of all malaria deaths in the region were children under 5 years old. Malaria has taken a major toll on human health for millennia, particularly for young children prior to the development of protective (though incomplete) immunity. Consequently, malaria has exerted a strong evolutionary selective pressure on the human genome [2]. Malaria can cause a wide spectrum of clinical disease, from asymptomatic parasitaemia to severe malaria syndromes and death. Understanding how human and parasite genetic variation interact with each-other and contribute to malaria disease severity is a key goal of malaria research.

Genetic variation at dozens of human loci have been proposed to provide protection against malaria [3–18]; although, some of these have not replicated in larger multi-centre studies [19,20]. It is likely that many host polymorphisms contribute to disease susceptibility, most with small effects [21]. Approximately 7% of the world’s population are thought to be carriers for inherited haemoglobin disorders, making them the most common human monogenic diseases [22]. Natural variation in the human haemoglobin genes was hypothesised to provide protection against malaria since at least the 1950s [15–17], given the central role of red blood cells (RBCs) in the *Plasmodium* life cycle, and the overlay of certain haemoglobin variants with malaria prevalence. However, the full scope of genetic diversity in human haemoglobin, and how this relates to malaria disease severity, is highly complex, and remains incompletely understood.

The dominant form of adult haemoglobin (HbA) comprises a tetramer of proteins transcribed from two α-globin genes and two β-globin genes (α_2_β_2_), responsible for oxygen transportation within RBCs. The α-globin and β-globin genes are located within the α- and β-globin loci on chromosomes 16 and 11, respectively. Of the haemoglobinopathies (altered haemoglobin makeup and/or structure), haemoglobin S (HbS, or sickle), haemoglobin C (HbC) and haemoglobin E (HbE) occur at significant population frequencies with distinct geographic distributions [23]. The sickle β-globin allele (β^S^) is a single point non-synonymous polymorphism changing Glutamic acid (coded by GTG) to Valine (GAG) at the sixth codon; this remains the strongest known association with protection against malaria in the human genome [19,20], maintaining high frequencies in populations exposed to malaria, primarily in Africa [24]. Homozygosity for β^S^ is the most common cause of sickle cell disease (SCD) in Africa. The central pathophysiology of SCD involves polymerisation of deoxygenated HbS, causing RBCs to become inflexible and misshapen, leading to haemolysis, endothelial dysfunction, inflammation and occlusion within the microcirculation [25]. Malaria protection is driven by the heterozygous state (HbAS) and fixation of the sickle allele cannot be attained due to the deleterious nature of the homozygote state (HbSS). The mechanisms by which heterozygous HbS provides malaria protection are still not fully understood, but several have been proposed (summarised in [26]).

HbS is present throughout sub-Saharan Africa to varying degrees [23,24], but a second haemoglobin type, HbC, occurs at high frequency only in central west Africa [27]. HbC arises due to an alternative single-point polymorphism in the sixth codon of β-globin (β^C^) at the base pair adjacent to the β^S^ polymorphism, leading to a switch from Glutamatic acid to Lysine. In west Africa, therefore, β-globin genotypes can include β^A/A^ (HbAA wild-type), β^A/S^ (HbAS sickle heterozygote), β^S/S^ (HbSS sickle homozygote), β^A/C^ (HbAC heterozygote), β^C/C^ (HbCC homozygote), and β^S/C^ (HbSC compound heterozygote). Many other polymorphisms in the β-globin gene have been described, including those that change gene expression causing β-thalassaemias [23,28], but these are less common in west Africa.

How variation in human and malaria parasite genomes interact with each-other and give rise to disease phenotypes remains an active area of research. A recent study of around 4,000 samples from children with severe malaria from Kenya and The Gambia identified three parasite loci that were associated with sickle haemoglobin, termed *P. falciparum* sickle-associated loci or *Pfsa1*, *Pfsa2* and *Pfsa3* [29]. *Pfsa1* and *Pfsa2* are on different loci on chromosome 2, while *Pfsa3* is located in a region of suspected structural variation on chromosome 11. The functional mechanism(s) underpinning these associations is unclear. Here, we have tested for associations between the human β-globin variants β^S^ and β^C^, and genome-wide variation in *P. falciparum* parasites, in a sample of mild malaria infections from a population in northern Ghana.

## Methods

### Study site and samples

Samples were collected from the Kassena-Nankana East Municipality and Kassena-Nankana West District, two adjoining administrative areas in the Upper East Region of northern Ghana, hereafter referred to as the KNDs. The population is primarily rural, and malaria transmission is seasonal, peaking during the rainy season from July to October. *P. falciparum* genetic diversity has been studied in this region previously [30] through a network of community health centres.

This study ran from 22/07/2015 to 30/08/2018, encompassing the rainy seasons for 2015, 2016 and 2017. Patients presenting to the Navrongo War Memorial Hospital (NWMH) and four satellite clinics in the KNDs with compatible symptoms for malaria were tested using an antigen Rapid Diagnostic Test (RDT) (CareStart™ malaria Pf (HRP2), Access Bio, NJ, USA). Participants were eligible if they tested positive for *P. falciparum* by RDT and informed consent was obtained and documented. Written consent forms were available in the predominant languages spoken in the region: Kasem, Nankam, and English. Four 50uL dried blood spots (DBS) and malaria blood smears for microscopy were collected from consenting participants. Parasitaemia was calculated by microscopy as asexual parasites per 200 white blood cells (WBC). DBS samples were transferred to the Wellcome Sanger Institute (WSI), UK, for DNA extraction and sequencing. DNA extraction from DBS was performed using the QIAamp Investigator Biorobot kit on the Qiagen Biorobot Universal instrument, protocol described in [31].

### Parasite whole genome sequencing and genotype calling

*P. falciparum* whole genome sequencing from DBS extractions was performed using the MalariaGEN pipeline established at the WSI, UK, described elsewhere [32]. Briefly, DBS samples undergo selective whole genome amplification (sWGA) to enrich for *P. falciparum* DNA [33], followed by short-read Illumina sequencing on the HiSeq platform. These samples are part of the broader MalariaGEN *P. falciparum* Community Project [32,34,35]. Genotype calling from the Illumina sequence data was performed at WSI using the MalariaGEN informatics pipeline [32]. All of the parasite genomes included in this study are available via the current iteration of the MalariaGEN *P. falciparum* Community Project, Pf7 [32].

### Human haemoglobin genotyping

Samples with ethical approval for use in this study (a subset of the MalariaGEN *P. falciparum* Community Project collection) were transferred via Beckman robot to new 96-well plate layouts organised by sample concentration (measured by Qubit High Sensitivity (HS) assay) and remaining volume after parasite sequencing. Human β-globin genotyping at codon 6 followed a PCR amplification and capillary sequencing strategy. Details on the primers and PCR conditions used can be found in the **Supplementary Methods**. The PCR products were capillary sequenced by the Eurofins company, using their PCR clean-up and Supreme Run service.

Capillary sequence chromatogram traces were visually inspected using the 4Peaks software tool (**Supplementary Methods**). The six callable β-globin genotypes from the sixth codon were: β^A/A^, β^A/S^, β^S/S^, β^S/C^, β^A/C^, and β^C/C^. Samples with ambiguous calls (eg. low amplitude chromatograph or chaotic uninterpretable signal) were repeat PCR-amplified and re-sequenced if available; if these samples remained ambiguous after the second attempt (or if there was no sample left for further attempts) then the sample was deemed to have failed β-globin genotyping. All β^A/S^, β^S/S^, β^S/C^, and β^C/C^ samples were repeat PCR-amplified and re-sequenced to confirm the result; there were no samples for which the second run disagreed with the first run if a call was successfully made on both runs.

### *P. falciparum* genome variant filtering for study population

Parasite genomic analyses were conducted using the data structures and curated sample metadata of the MalariaGEN Pf7 data resource [32]. Full details of variant filtering steps are described in **Supplementary Methods**. Briefly, the approximately 10 million variants in Pf7 were initially filtered to create a ‘working set’ of variants, including only biallelic SNPs that passed Pf7 Quality Control (QC) and had at least 0.1% allele frequency in the Ghana study population, yielding 123,845 SNPs. Heterozygous genotypes (representing mixed infections in *P. falciparum*) were regarded as missing. The previously reported lead associations in the *Pfsa* regions were explicitly included. This ‘working set’ of 123,845 SNPs was then further filtered for different analyses; the main logistic regression GWA analysis described in main text included 18,862 SNPs with a minor allele count of at least 10 among the 1,313 samples included in the analysis (after removing 55 closely related samples – described below). Genomic data manipulations from the Pf7 data resource were performed in Python; genotype and haplotype manipulations and downstream population genetic analyses were performed using the *scikit-allel* package.

### Population structure analysis

We used principal components analysis (PCA), as implemented in the QCTOOL package [36] to assess population structure in the 1,368 samples passing QC and with both parasite and β-globin genotypes in our data (further details in **Supplementary Methods**). For principal component computation, we restricted attention to the set of 14,352 SNPs (selected from the ‘working set’ of 123,845 SNPs) with at least 1% minor allele frequency and no more than 25% missing genotypes in these samples. To avoid genome-local effects, we further thinned this set based on genome coordinate so that no two SNPs in the thinned set were closer than 100bp apart, leaving 11,880 SNPs which we used for principal component (PC) calculation. Initial PCs (**Supplementary Figure 1**) revealed a small number of sample pairs with high relatedness values (estimated r > 0.5) that dominated the first two principal components. We therefore implemented a greedy algorithm to remove one of each sample pair having *r*_ij_ > 0.5. In total this removed 55 samples, leaving 1,313. We then re-computed principal components and genome-wide SNP loadings. Inspection of PCs and loadings suggested that the first two principal components correspond to potential population structure (indicated by reasonably similar loadings at SNPs across the genome) while subsequent PCs may reflect genome-localised or sample subset-specific effects (**Supplementary Figure 1)**. There was no evidence of population structure by sampling year, location, or HBB genotype (**Supplementary Figure 2**).

### Genome Wide Association

We performed a logistic regression analysis using the program HPTEST [29], which fits a logistic regression model with the host genotype as predictor and the parasite genotype as outcome, treating each variant separately (further details in **Supplementary Methods**). HPTEST reports a Bayes Factor reflecting the evidence for association as well as other summaries including an effect size estimate, standard error, and *P*-value. Interpretation of HPTEST results was previously discussed in detail [29]. For our main analysis, we used the 1,313 samples without closely related parasite samples identified through PCA, either with or without including the first two principal components. Data on sex was completely absent and data on age had high levels of missingness – age was recorded in 594/1555 (38%) of the samples that pass Pf7 QC. With such high missingness, age and sex were not used as covariates.

### Population genetics of the *Pfsa* loci

Allele frequencies and linkage disequilibrium (LD) for the *Pfsa* loci were computed in Python using the *scikit-allel* package. For the analysis of global patterns of *Pfsa* frequencies and LD, the total MalariaGEN Pf7 data resource (20,864 samples) was restricted to samples that passed Pf7 QC (N=16,203), and to countries with ≥100 QC pass samples, yielding 15,738 samples from 22 countries. One SNP was selected to represent each *Pfsa* locus for frequency and LD calculations: *Pfsa1* = chr2:631,190; *Pfsa2* = chr2:814,288; *Pfsa3* = chr11:1,058,035; *Pfsa4* (newly reported in this study) = chr4:1,121,472.

To analyse patterns of haplotype sharing at the *Pfsa* loci, we focussed on a 10kb section of the *P. falciparum* genome centred on the lead sickle-associated variant (identified as the variant with the highest Bayes factor) at each of the *Pfsa1-3* loci, using the MalariaGEN Pf7 data resource. At *Pfsa4*, we adjusted this to only include 7kb due to the higher density of variants at this locus. Ancestral vs. derived allele status at the *Pfsa* loci was determined as described in [37]. Further details on Pf7 variant filtering is described in **Supplementary Methods**. Genotypes were parsed into python code using the package *scikit-allel* (version 1.3.11) and clustered using the package *scipy* (version 1.14.1). Initially, the haplotype matrix was transposed so that each row represented a sample, and each column represented a variant. Hierarchical clustering was then performed using the average linkage method, which calculated the distance between clusters as the average distance between all pairs of genotypes in the clusters. The function used the resulting dendrogram to determine the order of the samples, reordered the haplotype matrix and the samples array accordingly, and returned the reordered haplotype matrix and samples array.

## Results

### Sample collection and β-globin genotypes

Blood samples were collected from people testing positive for *P. falciparum* with mild malaria who self-presented to community health centres in northern Ghana over a three year period, from 22^nd^ July 2015 to 30^th^ August 2018. *P. falciparum* whole genome sequencing was attempted on all 2,174 samples collected, as part of the MalariaGEN *P. falciparum* Community Project (Pf7 data release [32]). 1,555 (71.5%) samples successfully passed quality control (QC) filtering for the MalariaGEN Pf7 data resource. β-globin genotyping by PCR and capillary sequencing was attempted on all available samples following parasite sequencing, with 1,807 (83.1%) samples successfully assigned a β-globin genotype (**Figure 1**). 1,368 (62.9%) samples both passed Pf7 QC and had a β-globin genotype call and were used for genome-wide association (GWA) analysis, referred to as the study population.

**Figure 1.**
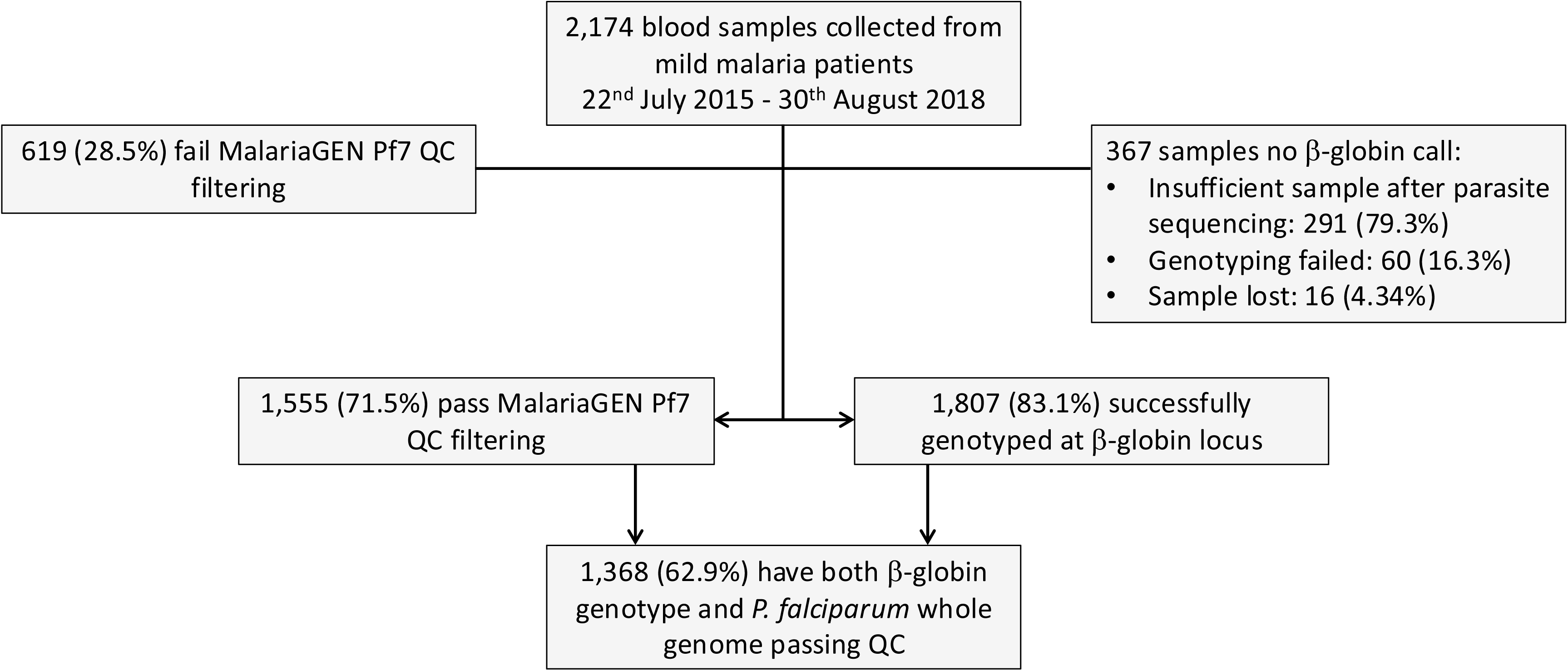
Study flow diagram.

Keeping the same nomenclature as that used by Band *et al*., the frequencies of *Pfsa1+* (chr2:631,190 T>A in the 3D7 v3 reference genome) and *Pfsa3+* (chr11:1,058,035 T>A) were 3.9% and 4.5% in the study population, respectively. The *Pfsa2+* variant (chr2:814,288 C>T) was not present in this population. At the human β-globin locus, β^A/A^ was the most common genotype identified in the study population (1008/1368, 74%), followed by β^A/C^ (312/1368, 23%) (**Table 1**). 36 samples (2.6%) included at least one β^S^ allele (β^A/S^, β^S/C^, or β^S/S^ genotypes), and 12 samples were β^C/C^.

**Table 1.**
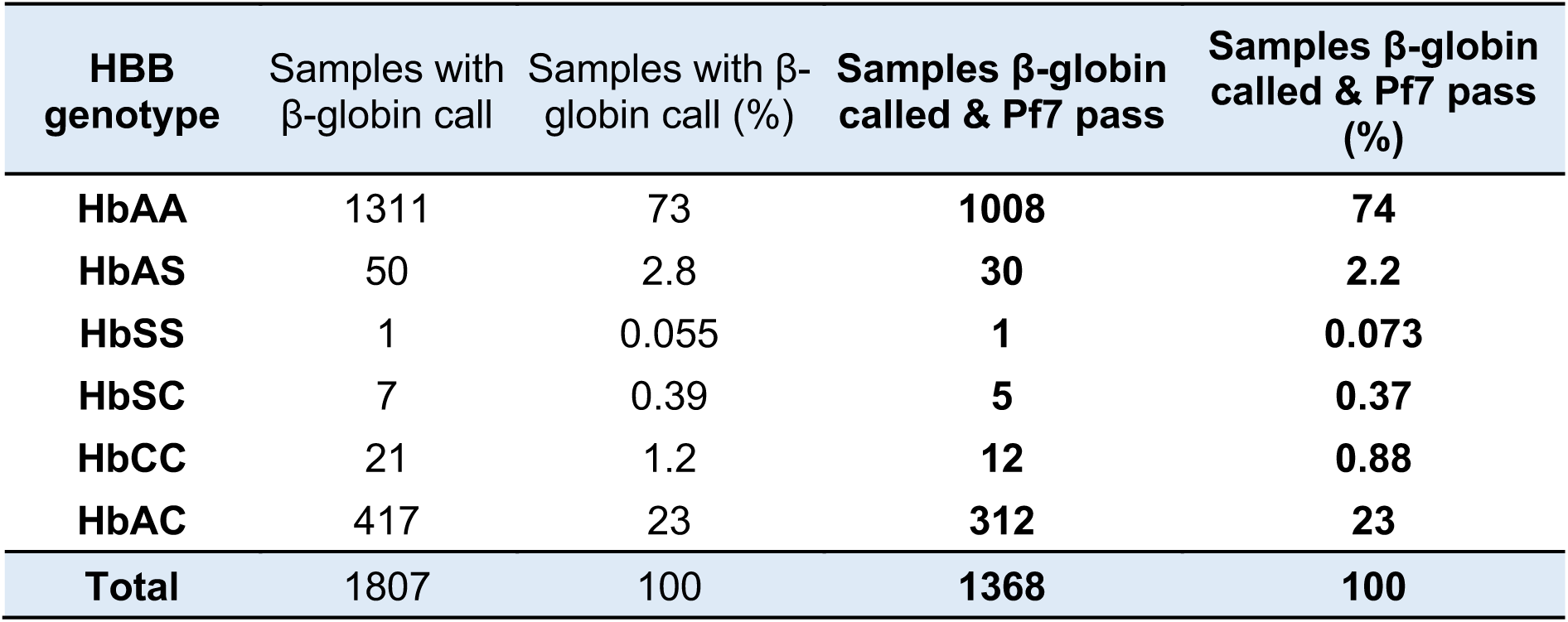
Counts of the different β-globin genotypes in all samples that had β-globin genotype calls available (N=1,807) and those that also had *P. falciparum* genomic data available that passed MalariaGEN Pf7 quality control filters (N=1,368). A full breakdown of sample counts by region, year and genotype is shown in **Supplementary Table 5.**

| **HBB genotype** | **Samples with β-globin call** | **Samples with β-globin call (%)** | **Samples β-globin called & Pf7 pass** | **Samples β-globin called & Pf7 pass (%)** |
| --- | --- | --- | --- | --- |
| **HbAA** | 1311 | 73 | **1008** | **74** |
| **HbAS** | 50 | 2.8 | **30** | **2.2** |
| **HbSS** | 1 | 0.055 | **1** | **0.073** |
| **HbSC** | 7 | 0.39 | **5** | **0.37** |
| **HbCC** | 21 | 1.2 | **12** | **0.88** |
| **HbAC** | 417 | 23 | **312** | **23** |
| **Total** | **1807** | **100** | **1368** | **100** |

### Associations between the *P. falciparum* genome and sickle haemoglobin

To investigate associations between *P. falciparum* variants and the human β-globin genotype, we applied a logistic regression approach using the human genotype as predictor and parasite genotype as outcome variable, after excluding a subset of 55 samples due to high relatedness (**Supplementary Methods** and **Supplementary Figure 1A**). Principal components analysis suggested some population structure was present (**Supplementary Figure 1B**) and we included two principal components as covariates to account for this. We used an indicator of β^S/*^ individuals (N=34; 33 β^A/S^ and 1 β^S/S^) versus β^no-S^ (N=1,279) as predictor, and tested for association with genotypes at each parasite variant in turn. Full results from this analysis can be found in **Supplementary Table 1** and **Figure 2**. This analysis revealed convincing replication of the previously identified associations at the *Pfsa1* locus (chr2: 631,190 T>A; BF_HbS_ = 5.5 × 10^7^, *P* = 1.3 × 10^−11^) and the *Pfsa3* locus (chr11:1,058,035 T>A, BF_HbS_ = 6.8 × 10^11^, *P* = 4.9 × 10^−16^). Observed effect sizes for these loci were comparable to those estimated previously in severe malaria cases (e.g. estimated odds ratio (OR) = 24, 95% confidence interval (CI) = 11 - 52 for the *Pfsa3* locus; compared to OR = 22, 95% CI = 10 - 47 across all data reported previously [29]). The *Pfsa2+* allele (chr2: 814,288 C>T) is rare in this population and was not tested.

**Figure 2.**
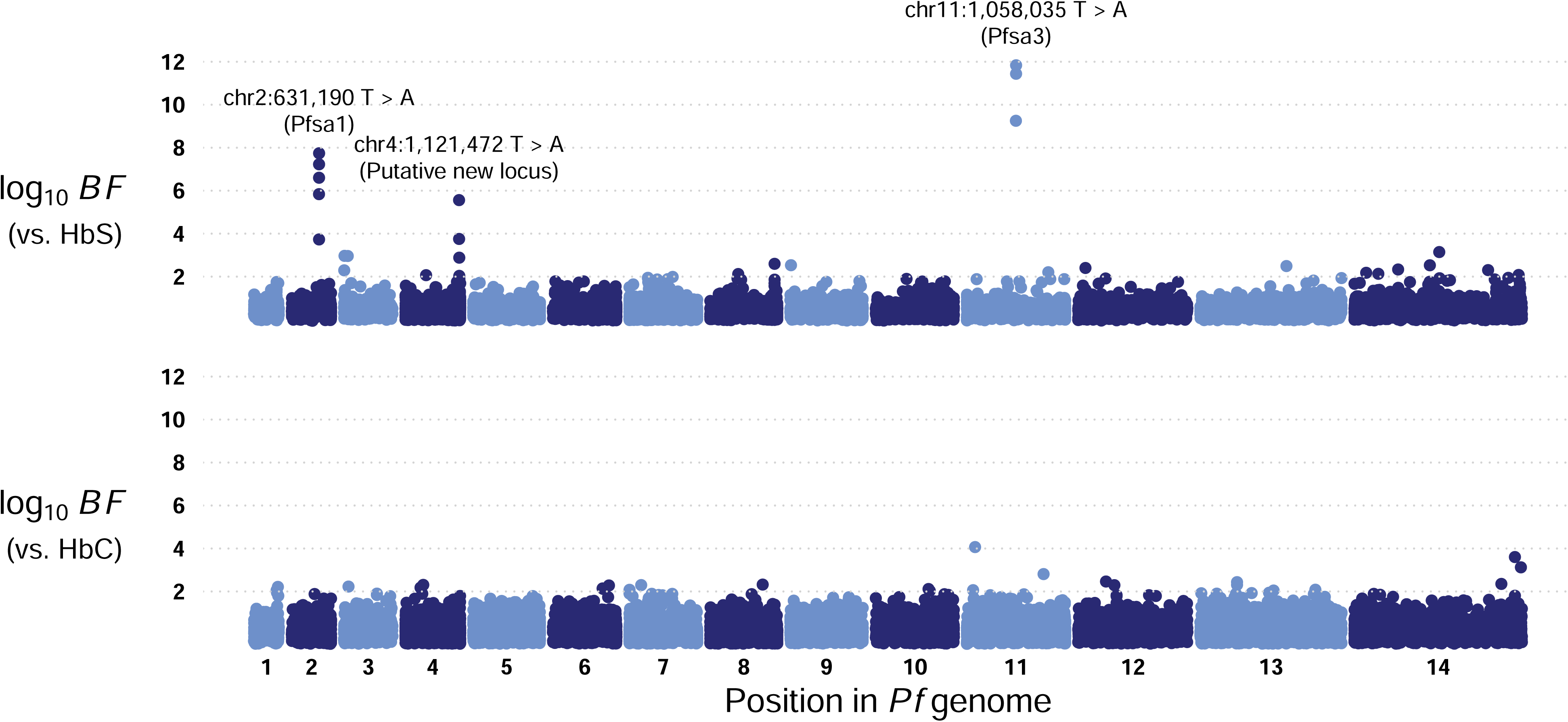
Evidence for association between haemoglobin variants and *P. falciparum* genetic variants in Ghanaian mild malaria cases. Plot shows the log10 Bayes factor for association with HbS (top row) and HbC (bottom row) (y axis), for SNPs across the parasite genome (x axis).

We observed evidence for association with a previously unreported locus on chromosome 4. The strongest evidence was seen at chr4:1,121,472 T>A (BF_HbS_ = 2.8 × 10^5^, *P* = 9.7 × 10^−9^; *OR* = 11.5, 95% CI = 5.0 - 26.5 for the ‘T’ allele) (**Figure 2**). This non-synonymous polymorphism occurs within the gene *FIKK4.2* (gene ID PF3D7_0424700 in the 3D7 reference), which has not previously been associated with sickle. (Of note, the 3D7 reference clone carries the associated ‘T’ allele, i.e. it is the ‘reference allele’ that was positively associated with sickle). A second nearby SNP (chr4:1,122,147 A>C) also had modest evidence for association (BF_HbS_ = 4.2 × 10^3^) (**Figure 3**). These results did not vary substantially when controlling for the first 5 PCs, or varying sample filtering criteria (e.g. BF_HbS_ = 5.3 × 10^5^; *OR* = 12.5 at chr4:1,121,472 when analysing a subset of 1,257 samples with a stricter relatedness criterion; Methods and **Supplementary Table 1**).

We attempted to replicate the putative new association by analysing two other datasets (**Figure 3**). The chr4:1,121,472 T>A SNP did not appear strongly associated in the previously reported set of severe malaria cases from The Gambia [29] (estimated OR = 1.68 for the ‘T’ allele, 95% CI = 0.61 - 4.64); an accurate estimate was not possible in Kenya where the T allele is at extremely low frequency. However, we observed convincing replication of this association in 32 previously reported mild malaria cases from Mali, for which RNA-seq data has previously been reported [38]. Excluding 5 samples which appeared to have mixed bases in RNA reads at this locus (requiring < 5% or > 95% of reads at the locus to support expression of the T allele), we found strong evidence for replication: 11 of 14 (79%) infections of HbAS children carried the associated ‘T’ allele; while 11 of 13 (85%) HbAA infections carried the alternative ‘A’ allele; OR = 20, 95% CI = 2.8-145; *P*=2.8×10^−3^. We saw no evidence in these samples that total read counts (reflecting expression levels) were associated with β^S^ status or with the chr4:1,121,472 T>A genotype.

**Figure 3.**
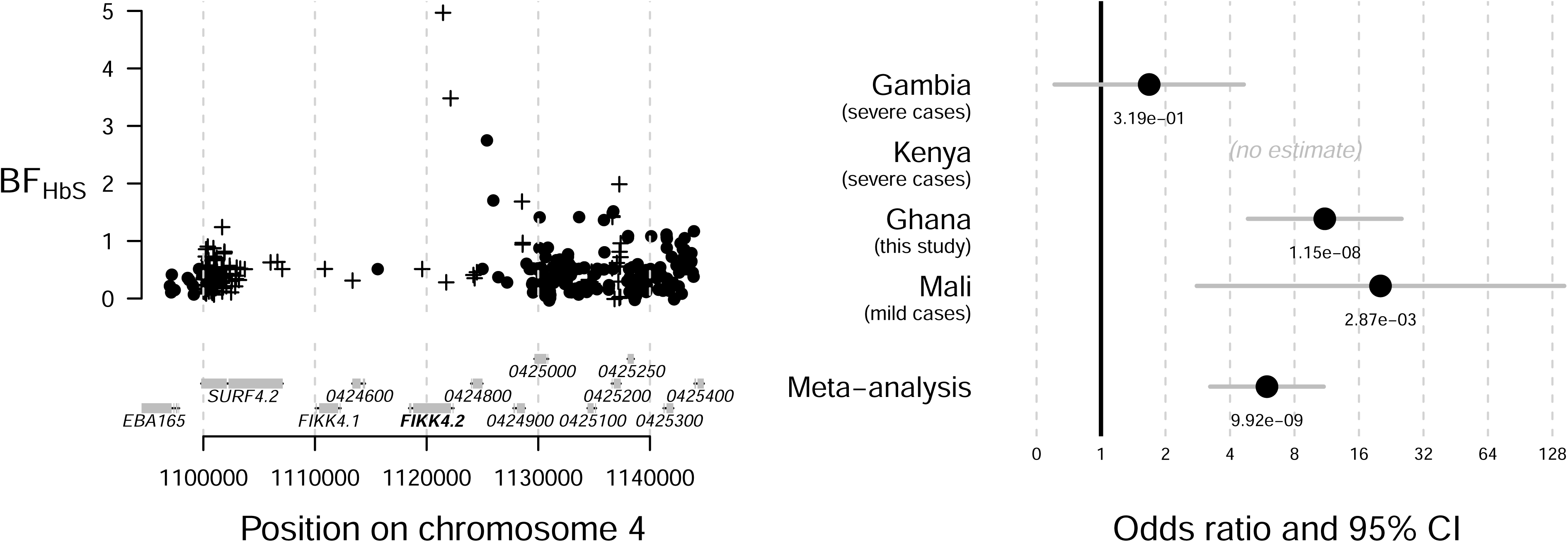
Discovery and replication evidence for association between HbS and *Pfsa4*. The left hand side plot depicts the log10 Bayes factor for association with HbS (y axis) for SNPs in the parasite genome zoomed onto the *Pfsa4* locus (x axis). The top hit is SNP chr4:1,122,147, ‘T’ allele. The right hand plot shows point estimates with 95% confidence intervals for the Pfsa4-HbS association, estimated using logistic regression from across multiple studies. The populations are: Gambian severe cases from [29], Kenyan severe cases from [29], Ghanaian cases from this study, and Malian cases from [38]. Text indicates the 2-sided *P*-value for each point.

**Figure 4.**
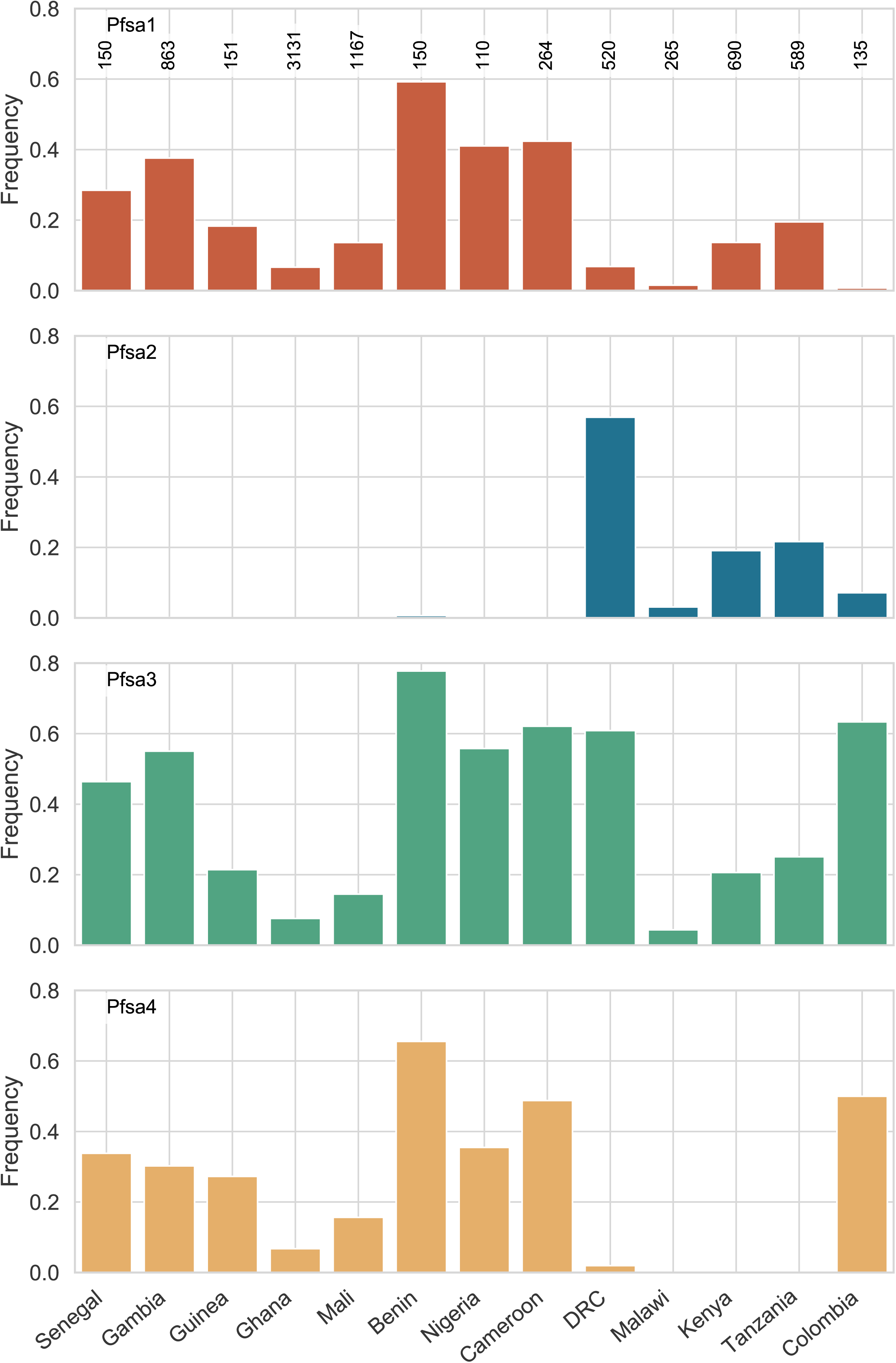
Global distribution of *Pfsa1-4+* alleles in the MalariaGEN Pf7 data resource. Data from 15,738 samples from 22 countries, which passed Pf7 QC filters and requiring ≥100 samples for each country. Countries are organised left to right approximately from West to East Africa, with Colombia as the only non-African country represented. The *Pfsa*+ alleles were very low or absent from all Asian countries in Pf7, not shown on Figure.

Taken together, these results provide suggestive evidence that the chr4:1,121,472 T>A SNP is associated with β^S^ in mild malaria infections in west Africa, but raise questions as to why the association is not also seen in the severe malaria data from Kenya and The Gambia analysed previously, discussed further below. Following previous work, we tentatively refer to this new locus as *P. falciparum* sickle-associated locus 4 (*Pfsa4*), and to alleles that are associated or not associated with sickle as *Pfsa4+* (‘T’ allele, present in the 3D7 reference genome) and *Pfsa4-* (‘A’ allele, alternative to the 3D7 reference genome), respectively. In this Ghanaian cohort of mild malaria cases, the *Pfsa1+*, *Pfsa3+*, and *Pfsa4+* allele frequencies differed markedly between sickle and non-sickle individuals (**Supplementary Figure 3**). For example, *Pfsa4+* was present in 12/36 (33%) of individuals carrying at least one sickle allele, compared with 3% of people with HbAA (**Supplementary Table 3**). Individuals carrying at least one sickle allele were around 10x more likely to be infected by parasites carrying at least one *Pfsa*+ allele than by parasites that did not carry any of these alleles (50% vs 5%, respectively). Thus, despite the prevalence of sickle carriage among this Ghanaian cohort being approximately 3%, and the prevalence of *Pfsa1*+*, Pfsa3+* and *Pfsa4+* alleles in *P. falciparum* being around 4%, over half of the malaria infections encountered by sickle individuals were caused by parasites carrying at least one *Pfsa*+ allele.

Because notable linkage disequilibrium (LD) exists between the putative *Pfsa4* locus and the previously identified *Pfsa1* and *Pfsa3* loci (discussed further below), there is the possibility that the association at *Pfsa4* is driven by LD to truly associated signals and does not represent an independent signal. To assess this, we reperformed the association analysis within the Ghanaian, and previously published Gambian and Kenyan datasets, using genotypes at the *Pfsa1* and *Pfsa3* loci as covariates (**Supplementary Figure 4**). The effect of HbS on *Pfsa4* genotype is almost completely diminished by the inclusion of *Pfsa1* or *Pfsa3* as covariates in Ghana. This might suggest that *Pfsa4* does not represent an independent association to HbS. However, inspecting this relationship at the other identified *Pfsa* loci, we observe that the signals of association at the *Pfsa1*, *Pfsa2* and *Pfsa3* loci also diminish to a similar degree when conditioned on *Pfsa1* and *Pfsa3* genotype within severe cases in Kenya. This is not the case in severe cases from The Gambia, however, where the HbS-*Pfsa* association is largely unchanged by the inclusion of *Pfsa* genotypes as covariates. This is likely to be explained by lower levels of LD between the *Pfsa1* and *Pfsa3* loci within The Gambia compared to within Kenya and Ghana. Therefore, although HbS-*Pfsa* associations do not appear statistically independent of one another, they are connected through prominent LD, despite being situated far apart in the *P. falciparum* genome across multiple chromosomes. This is challenging to explain without a biological relationship between them and HbS.

To explore whether the effects of *Pfsa* loci were additive, we computed the Ratio of Relative Risk (RRR) for different combinations of *Pfsa* genotypes occurring in HbS-carrying individuals (**Supplementary Methods**). The RRR increased in infections carrying *Pfsa1*+, *Pfsa3*+ and *Pfsa4*+ alleles combined, compared to those carrying only *Pfsa1*+ and *Pfsa3*+. This analysis suggests that *Pfsa4+* does have an effect in improving the ability of *P. falciparum* parasites to infect HbS individuals, compared to *Pfsa1+* and *Pfsa3+* alone (**Supplementary Figure 5**).

### Examining the putative *Pfsa4* locus

*FIKK4.2* is a member of a larger family of FIKK serine/ threonine kinase proteins. It is exported to the erythrocyte cytosol during blood-stage infections, and, along with other FIKKs, is thought to regulate phosphorylation of erythrocyte and parasite proteins in the erythrocyte membrane. FIKK4.2 has also been associated with specific compartments in the infected red cell (termed K-dots [39]). In the 3D7 genome assembly, chr4:1,121,472 lies in the second exon of the *FIKK4.2* gene, and encodes a Leucine → Isoleucine change at amino acid 961. This is within the kinase domain, and is downstream of a lengthy segment of low complexity repeat sequences [39].

Specifically, an amino acid hexamer repeat occupies a significant proportion of FIKK4.2 and is located nearby to the top candidate SNP. The majority of repeats take the form SD[HNS]NH[KM]. We undertook a preliminary analysis of this region using multiple sequence alignment of 16 high-quality *P. falciparum* reference genomes [40] (**Supplementary Methods**). FIKK4.2 varied in length between these isolates from 1,079 amino acids (isolate PfKH01) to 1,223 amino acids (isolate Pf3D7). This variation reflected expansion / contraction of the repeat sequence, which varied from 402 to 546 amino acids (67 - 91 hexamer repeats). Notably, the Pf3D7 sequence, with 91 repeats, was substantially longer than that in any other assembly; the next longest being Pf7G8 (protein length 1,181; 84 hexamer repeats). The major cause of this increased length was a triplication of a DNA sequence in Pf3D7 of approximately 170bp, at positions 1,120,534-1,120,005; this repeat was only present in one or two copies in each of the other 15 genome assemblies (**Supplementary Figure 6**). In 3D7, this repeat unit ends at position 1,121,413, which is only 59 bases away from the lead *Pfsa4* SNP. Intriguingly, the *Pfsa4+* allele is present in 4/16 reference genomes, all of which contain two or three copies of the repeat region; whereas of the 12/16 isolates with the non-associated allele at *Pfsa4*, 11/12 only have a single copy of the repeat section (*P*<0.05 for 11/12 isolates having one copy, if expectation was for 12/12 to have two or three copies as per *Pfsa4+* isolates, exact Binomial test). This raises the possibility that structural variation within the hexamer repeat segment of FIKK4.2 is causally involved in the sickle association signal, and the *Pfsa4+* SNP could be a bystander. However, as this finding is based on 15 genomes, it should be considered suggestive at this stage pending investigation with a larger sample size.

### Little evidence of association with HbC

One potential explanation for effects specific to central west African populations, could be the presence of the haemoglobin C variant in these populations. We conducted a second GWA analysis to test for association between parasite variants and individuals possessing at least one HbC allele. To avoid confounding by the HbS association we excluded all individuals carrying HbS alleles, leading to an analysis (after removing highly related individuals) of β^C/no-S^ (N=315; 303 β^A/C^ and 12 β^C/C^), vs β^A/A^ individuals (N=964). Applying the same methodology as with the sickle analysis, we performed logistic regression using HPTEST. In contrast to HbS, we did not identify any strong signals of association with HbC (**Figure 2, Supplementary Table 2**). The strongest association observed was seen at chr11:269,007 T>C (BF = 1.2×10^4; P = 0.00016) which lies in a conserved protein of unknown function (PF3D7_1106500); however, this SNP was relatively rare (∼1% frequency). Our study was adequately powered to detect HbC-associations in the *P. falciparum* genome of a similar effect size to the HbS-*Pfsa* associations for relatively common SNPs (5% frequency or above) (**Supplementary Methods**), but will be underpowered for rarer variants or where the effect sizes are smaller.

### Population genetics of the *Pfsa* loci

We investigated linkage disequilibrium (LD) between the *Pfsa* loci. As expected, nearby SNPs from the *Pfsa3* locus on chromosome 11 (chr11:1,058,035 T>A, chr11:1,057,437 T>C, and chr11:1,054,282 T>C) were all in high LD with each-other (*r^2^* 0.76 – 0.95), as were the four SNPs identified at the *Pfsa1* locus on chromosome 2 (chr2:630,545 C>T, chr2:631,190 T>A, chr2:630,290 A>T, and chr2:629,375 C>G; *r^2^* 0.91 – 0.98). The *Pfsa4* SNP with strongest evidence of association with sickle (chr4:1,121,472 T>A) was in LD with the nearby SNP identified in the GWA analyses (chr4:1,122,147 A>C, *r^2^* = 0.84). As noted by Band *et al*., *Pfsa1* and *Pfsa3* loci were in high LD with each-other despite being located on different chromosomes. For example, the top hit SNPs from the *Pfsa1* and *Pfsa3* loci had an *r^2^* of 0.62. *Pfsa4*, on chromosome 4, was also in relatively high LD with both *Pfsa1* and *Pfsa3* loci. For example, the top hit SNP at *Pfsa4* (chr4:1,121,472 T>A) had an *r*^2^ of 0.22 and 0.24 for the top hit SNPs at the *Pfsa1* and *Pfsa3* loci, respectively. These data suggest that *Pfsa1+*, *Pfsa3+* and *Pfsa4+* alleles are maintained together despite being located on different chromosomes, consistent with having linked biological functions.

Next, we investigated the global distribution of *Pfsa1-4* alleles using the MalariaGEN Pf7 data resource [32]. Of the 20,864 samples in Pf7, we limited the analysis to samples that passed MalariaGEN Pf7 QC (N=16,203), and to countries with ≥100 QC pass samples, yielding an analysis set of 15,738 samples from 22 countries. The *Pfsa1-4+* alleles were almost completely absent from Asia, but were present at varying frequencies in Africa and the only South American country represented, Colombia (**Figure 5**). *Pfsa1+* and *Pfsa3+* were distributed widely across Africa but with substantial heterogeneity in prevalence; for example, the frequency of *Pfsa1+* in Benin was 59%, compared with 2% in Malawi (**Supplementary Table 3**). *Pfsa4+* was essentially absent from east African populations, and was at very low frequency in the Democratic Republic of the Congo (DRC), but was found at much higher frequencies in west African populations (7% - 66%). The inverse pattern was seen in *Pfsa2+*, which was prevalent in east Africa and the DRC but almost completely absent from west Africa. Intriguingly, *Pfsa3+* and *Pfsa4+* were also present at high frequencies (63% and 50%, respectively), and *Pfsa2+* at lower frequency (7%), in Colombia.

**Figure 5.**
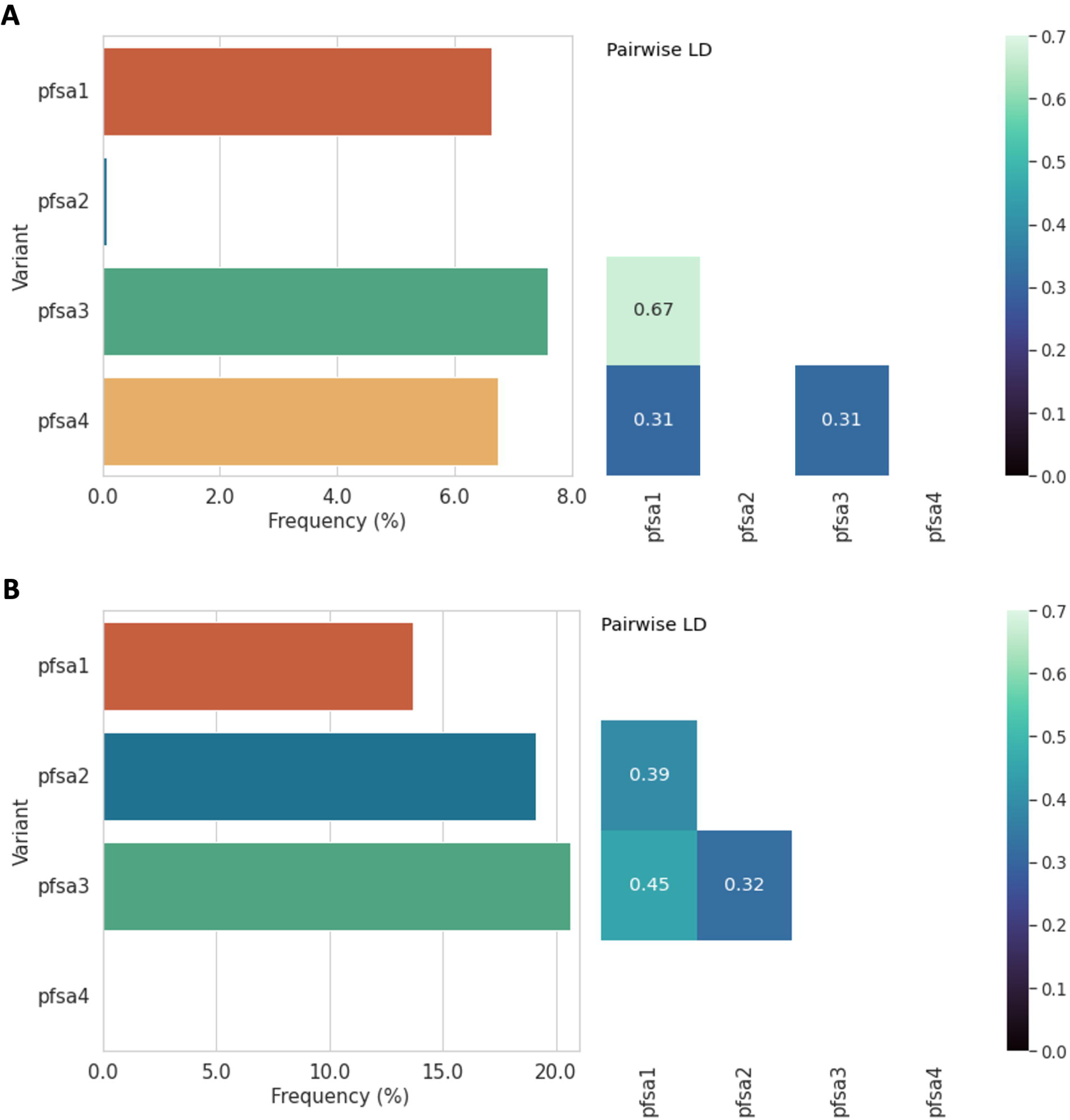
Frequencies and linkage disequilibrium (LD) of *Pfsa1-4+* alleles in Ghana (top) and Kenya (bottom). These well-sampled countries are shown to illustrate the distinct patterns of *Pfsa1-4* population genetics observed in different African countries.

There was evidence of haplotype sharing at each of the *Pfsa1-4* loci in parasites carrying the sickle-associated alleles in the MalariaGEN Pf7 data resource, consistent with each *Pfsa+* allele having a common evolutionary origin at each locus (**Supplementary Methods, Supplementary Figure 7**). However, we did not find evidence that any of the *Pfsa1-4* loci were under strong recent positive selection, based on integrated haplotype scores (iHS) (**Supplementary Table 4**, **Supplementary Methods**).

Globally, in the Pf7 dataset, patterns of LD were complex and varied between populations (illustrated by Ghana and Kenya in **Figure 5**; all countries shown in **Supplementary Figure 8**). In west Africa, where the combination of *Pfsa1+*, *Pfsa3+* and *Pfsa4+* was present, SNPs at these loci had relatively high LD with each-other in Guinea, Ghana and Mali, but weaker LD in Senegal and The Gambia. In east Africa, where the combination of *Pfsa1+*, *Pfsa2+*, and *Pfsa3+* is present, these loci were in high LD with each-other in Malawi, Kenya and Tanzania; but LD was lower for these loci in the DRC. In Colombia, *Pfsa3+* and *Pfsa4+* had high LD (*r*^2^= 0.49).

In summary, these data reveal complex patterns of *Pfsa1-4+* allele frequencies and associations between populations. *Pfsa1+* and *Pfsa3+* are present across Africa, *Pfsa2+* is more localised to east Africa, and *Pfsa4+* more localised to west Africa; within these regions, the extent to which *Pfsa+* alleles correlate with each-other is variable, ranging from very high (e.g. *r*^2^ of 0.67 for *Pfsa1+* and *Pfsa3+* in Ghana), to zero (e.g. *Pfsa3+* and *Pfsa4+* in Benin).

## Discussion

We have investigated host-parasite interaction by searching for associations between variation in the *P. falciparum* genome with human variation in the β-globin gene in a population with mild malaria from northern Ghana. Two parasite loci previously associated with sickle haemoglobin in severe malaria cases from Kenya and The Gambia [29] were replicated in this cohort, termed *Pfsa1* and *Pfsa3*. A putative new locus of association on *P. falciparum* chromosome 4 was identified within the gene *FIKK4.2*, which we name *P. falciparum* sickle associated locus 4, or *Pfsa4*. Further work is needed to validate this locus, both experimentation to determine its mechanistic function and increasing the sample size of sickle individuals being studied, as this study was limited to only 36 HbS-carrying individuals. The signal at this potential new locus was not associated with sickle in the severe malaria cases from The Gambia, and the variant is absent in Kenya. However, the *Pfsa4+* association with sickle did replicate in a sample of mild malaria cases from Mali. Therefore, evidence for the Pfsa4 candidate signal was only identified in West Africa and its broader generalisability remains to be established with additional datasets. There were no significant associations found between parasite SNPs and β^C^ genotypes. The geographic distributions of the *Pfsa1-4* alleles vary with distinct patterns observed in western and eastern Africa, explored further in [37]. Haplotype sharing indicates a common evolutionary origin for each *Pfsa* locus. The lack of strong positive selection at these loci, and their broad geographic distributions, suggests they have not emerged in the very recent evolutionary past.

The high prevalence of β^C^ and relatively low prevalence of β^S^ observed in this study is broadly consistent with previous results. HbC has mainly been described in west Africa, and it has been hypothesised that the relatively more benign phenotype of HbCC compared with HbSS would lead to the gradual replacement of the β^S^ allele by β^C^ in west Africa [13]. The proportion of β^S^ carriers we observed was lower than other studies in the region [41,42]. Around 2% (15,000) of Ghanaian newborns are diagnosed with sickle cell disease (SCD) annually [43,44]; the majority of patients with SCD attending a large teaching hospital in Accra were found to have HbSS (55.7%) or HbSC (39.6%) genotypes [43]. The relatively lower prevalence of β^S^ alleles we observed may be partly because sickle-containing genotypes protect against malaria disease, so the prevalence of such genotypes in this population, who were selected for having mild malaria, would be lower than in an unselected population. Variation in HbS prevalence may also be due to resource limitations in available healthcare to support children born with SCD in rural populations versus urban centres in Accra.

The replication of the association between *Pfsa1+* and *Pfsa3+* alleles with sickle in people with mild malaria is consistent with the view that parasites possessing these variants can bypass the protective effects of sickle and cause both mild and severe malaria syndromes in sickle individuals more readily than parasites lacking these variants. On this view, the sickle-associated variants effectively open an ecological niche (individuals carrying β^S^ alleles) that is otherwise relatively closed off (or restricted) for *Pfsa*-parasites; rather than the view that these variants increase severity in sickle individuals *per se*. The age of the β^S^ mutation is uncertain, with estimates suggesting a single origin around 20,000 years ago (range: 13,000-70,000 years [45]). Selection for *Pfsa*+ mutations may have occurred once β^S^ reached appreciable frequencies. It is currently unclear what has prevented *Pfsa*+ parasites from outcompeting *Pfsa*-parasites in regions where HbS is common but geographical variation may play a role [37]. The frequency of *Pfsa*+ polymorphisms may depend on the regional prevalence of HbS, and the relative fitness advantage of *Pfsa*+ alleles in HbS RBCs versus their disadvantage in HbA RBCs, compared with *Pfsa*-alleles. Moreover, sexual recombination during the mosquito phase would be expected to mix *Pfsa*+ and *Pfsa*-alleles. The presence of high LD between the *Pfsa*+ polymorphisms is consistent with an epistatic effect on parasite fitness counter-acting the effect of sexual recombination, to maintain the *Pfsa*+ alleles together. It is unclear whether all of the *Pfsa*+ alleles play a direct and independent role in assisting parasites to survive in HbS-carrying hosts, or whether some only mediate their effect in the context of other *Pfsa* alleles, for example to compensate for fitness costs. Further experimental work is required to explore these effects functionally.

The putative *Pfsa4* locus was found within the gene encoding the serine/ threonine kinase FIKK4.2. The FIKK kinase family (characterised by a phenylalanine (F) – isoleucine (I) – lysine (K) – lysine (K) motif) has expanded in the *Laverania* clade (which includes *P. falciparum* and other *Plasmodium* species that infect great apes), comprising around 19 proteins in *P. falciparum* distributed over 11 chromosomes. The majority of *P. falciparum* FIKK kinases are exported into the host Red Blood Cell (RBC), and each is associated with a unique ‘phosphorylation fingerprint’ of host RBC proteins [46]. There is evidence that FIKK kinases play a role in remodelling the RBC surface and altering the mechanical properties of infected RBCs [47,48]. RBCs infected by parasites lacking FIKK4.2 through experimental knockout were less rigid and less adhesive compared with RBCs infected by wild-type parasites despite unchanged levels of *P. falciparum* Erythrocyte Protein 1 (PfEMP1) expression, suggesting a role in malaria pathogenesis [39]. FIKK kinases have been proposed as potential targets for antimalarial drug design [49]. Intriguingly, the lead *Pfsa4+* SNP identified in our GWA analysis sits very close to a lengthy repeat region. Analysis of 16 high-quality *P. falciparum* genome assemblies tentatively suggests that the *Pfsa4+* SNP may be associated with increased copy number in this repeat section. Calling copy number variation (CNV) from short read sequencing data following parasite selective whole genome amplification (sWGA) [50] is challenging, so further work is required to assess this locus at population scale and to investigate functional effects of FIKK4.2 variants including CNV. In summary, the Pfsa4 locus can be considered a candidate signal of association with sickle, in high LD with the originally identified Pfsa1 and Pfsa3 loci, rather than a definitively confirmed, independent locus, pending further evidence for association and mechanistic experiments.

The lack of association between β^C^ alleles and parasite variation may reflect the fact that the vast majority of individuals possessing a β^C^ allele were genotype β^A/C^, which is thought to confer a reduced protective effect against *P. falciparum* relative to β^C/C^ [13], and likely also less than β^A/S^; the strength of selection acting on the parasites by β^A/C^ individuals will therefore be accordingly weaker. With only 12 β^C/C^ individuals in our cohort, the sample size is too small to analyse this sub-group. The effect specifically of the β^S/C^ genotype on malaria protection and *P. falciparum* genetic associations is, to our knowledge, unknown, as is the effect of other compound heterozygotes such as β^S^ with thalassaemia.

This study has several limitations. The small number of individuals carrying β^C^ and β^S^ alleles significantly limits the statistical power of the GWA analysis, meaning only variants with large effect sizes could be detected. Associations with smaller effect sizes and/or specific to certain β-globin genotypes such as β^C/C^ will require larger sample sizes for discovery. Further work is needed to confirm that the potential new locus of association replicates in other, larger population cohorts with more HbS genotypes (and other haemoglobin variants) represented. Our study does not distinguish whether all of the *Pfsa* loci are directly causally associated with sickle haemoglobin, or whether some of these loci are associated with each-other (i.e. in linkage) but not directly interacting with HbS. However, the fact these loci, located on different chromosomes, are in high LD with each-other is suggestive of some linked functional role, even if indirectly connected to HbS. Functional experimentation is required to unravel the mechanistic role of each *Pfsa* allele and how they are interacting in the infected RBC. This study extends previous work that focused on severe malaria by recruiting mild malaria cases; future work can explore parasite-sickle associations in asymptomatic individuals. The method of genotyping the human β-globin locus involved PCR amplification and capillary sequencing a small region at the start of the β-globin gene and did not account for other forms of genetic diversity in the haemoglobin genes. For example, it is possible that some of the homozygous calls were in fact compound heterozygotes with a β-thalassaemia allele. Although, complete β-globin gene deletion is an uncommon mechanism of β-thalassaemia [51], and β-thalassaemia is less common in west Africa than in other populations [23]. Minimal population demographic data were available to control for potential confounders, such as participant ethnic group. However, the logistic regression analysis was performed after removing samples that appeared highly similar, and incorporated principal components as covariates to account for population structure. This study does not have a matched control group who did not have clinical malaria against which to compare the β-globin genotype frequencies. This study therefore cannot directly assess the extent to which *Pfsa+* alleles reduce the protective effect of sickle against mild malaria relative to people without symptomatic malaria. However, their association with β^S^ alleles in mild malaria, combined with the fact that the same variants reduce the protective effect of sickle against severe malaria relative to population controls [29], supports the view that these variants allow parasites to access sickle individuals which wild-type parasites otherwise struggle to infect, and therefore reduce the protective effect of sickle against both mild and severe malaria. The GWA analysis was limited to high-confidence SNPs; CNV and other more complex forms of genetic variation were not included, due to the risk of spurious calls from short read post-sWGA data. Future work can explore these challenging forms of genetic variation further, potentially leveraging long read sequencing platforms and functional experimentation. This is particularly relevant given the potential link between CNV and the lead *Pfsa4+* SNP identified in our study.

## Conclusions

In conclusion, this study adds further evidence that several loci in the *P. falciparum* genome are associated with sickle haemoglobin, in both mild and severe malaria. Variants at these loci are maintained in linkage with each-other despite being on separate chromosomes. This suggests that in a population, subsets of *P. falciparum* parasites will inherit genetic traits that make them better suited to particular human genetic contexts, such as ‘specialists’ at infecting sickle RBCs. The frequencies of different host genotypes, and the relative fitness costs of the polymorphisms for the parasite in non-sickle hosts, will sculpt the evolutionary pressures acting on these loci in the parasite, which may in turn affect the selection pressures acting on the human genome in an ancient coevolutionary process. Future work could validate and extend these results by recruiting a larger population to ensure robust statistical power to analyse different haemoglobin genotypes, incorporate more genetic variation from both human and *P. falciparum* genomes, and establish functional data to uncover the mechanisms of how these variants help the parasite to infect people carrying sickle haemoglobin alleles.

## List of abbreviations

CNV: Copy number variation
FIKK: Phenylalanine (F) – isoleucine (I) – lysine (K) – lysine (K) motif
GWA(S): Genome wide association (study)
HbA: Haemoglobin A
HbC: Haemoglobin C
HbE: Haemoglobin E
HbS: Haemoglobin S
LD: Linkage disequilibrium
PCA: Principal components analysis
PCR: Polymerase chain reaction
PfEMP1: P. falciparum Erythrocyte Protein 1
Pfsa: P. falciparum sickle-associated locus
QC: Quality Control
RBC: Red blood cell
RDT: Rapid Diagnostic Test
RRR: Ratio of Relative Risk
SCD: Sickle cell disease
SNP: Single nucleotide polymorphism
sWGA: Selective whole genome amplification
WHO: World Health Organization
WSI: Wellcome Sanger Institute

## Declarations

### Ethics approval and consent to participate

Ethical approval for the study was granted by the Navrongo Health Research Centre Institutional Review Board (NHRCIRB203). All participants or their guardians (as appropriate) were provided detailed information sheets and gave informed consent prior to enrolment. Further approval was granted by the Wellcome Sanger Institute’s Research Ethics Committee for the analysis of the samples. Sample collection was conducted by LNA, who anonymised the samples prior to making them available to the Wellcome Sanger Institute. All patient-identifiable data are securely stored by LNA and only non-patient identifiable were provided to the Wellcome Sanger Institute. The study complies with all relevant ethical regulations.

This research was conducted collaboratively between researchers based at the University of Ghana, the Navrongo Health Research Centre (NHRC), the Wellcome Sanger Institute, and University of Oxford. Study conceptualisation, data analysis and interpretation, and authorship of publications were shared jointly between personnel based in Ghana and the UK.

### Availability of data and materials

The *P. falciparum* variant calls (VCF file) are available to download open-access via the MalariaGEN *P. falciparum* Community Project Pf7 resource. (The VCF file contains variant calls for all samples in Pf7, which includes all of the samples analysed in this study). The β-globin genotype calls linked to Pf7 sample codes are available in **Supplementary Table 6**. The filtered parasite genotype calls derived from Pf7 and associated human beta-globin genotype calls can also be downloaded from FigShare, details below:

Hamilton, William (2025). Analysis datasets used for Pfsa4 genome-wide association analysis (Hamilton et al).. figshare. Dataset. https://doi.org/10.6084/m9.figshare.30705047

Source code for QCTOOL and HPTEST is available at: code.enkre.net/qctool, and on Zenodo (doi:10.5281/zenodo.5685581). The code for running the GWAS is available via GitHub, link: https://github.com/wlhamilton/pfsa4/tree/main.

### Authors’ contributions

Conceptualization: LNA, WLH, DPK

Investigation: WLH, AJF, ED, GB, LNA

Formal analysis: WLH, AJF, GB, LNA

Resources: KAR

Data Curation: WLH, LNA

Writing – original draft preparation: WLH, GB, LNA

Writing – review and editing: WLH, GB, LNA, KAR, YA, VA

Visualization: WLH, AJF, GB

Supervision: LNA, GB, DPK

Project administration: LNA, YA, VA

Funding acquisition: DPK, LNA, GAA

## Acknowledgements

We thank the teams in the Navrongo Health Research Centre (NHRC) and the Wellcome Sanger Institute (WSI) who generated the *P. falciparum* genome sequence data as part of the MalariaGEN *P. falciparum* Community Project, and the study participants and their guardians who provided the samples. Thank you to Richard Pearson for his help accessing and using the Pf7 data resource, and to Christen Smith, Sonia Goncalves and Rachel Ogidan for support ensuring legal and ethical compliance. Thank you to Kevin Esoh for contributions to discussions of *P. falciparum* population genetics. This work is dedicated to the memory of Professor Dominic Kwiatkowski, who died prior to its completion.

