## Supplementary Materials for "Associations between the *Plasmodium falciparum* genome and sickle haemoglobin identified in mild malaria cases from Ghana"

**Authors**

**William L. Hamilton[1,2,3 *], Annie J. Forster[4], Yaw Aniweh[5], Victor Asoala[6], Eleanor Drury[1], Kirk A. Rockett[4], Gordon A. Awandare[5], Dominic P. Kwiatkowski[1], Gavin Band[4], Lucas N. Amenga-Etego[5 *]**

**Author affiliations**

1. Wellcome Sanger Institute, Wellcome Trust Genome Campus, Hinxton, CB10 1RQ, United Kingdom
2. University of Cambridge, Department of Medicine, Cambridge Biomedical Campus, Hills Road, Cambridge CB2 0QQ, United Kingdom
3. Cambridge University Hospitals NHS Foundation Trust, Cambridge Biomedical Campus, Hills Road, Cambridge CB2 0QQ, United Kingdom
4. Centre for Human Genetics, Roosevelt Dr, Headington, Oxford OX3 7BN, United Kingdom
5. West African Centre for Cell Biology of Infectious Pathogens (WACCBIP), University of Ghana, Legon, Accra, Ghana
6. Navrongo Health Research Centre (NHRC), Ghana Health Service, Navrongo, Upper East Region, Ghana

*** Corresponding authors**

Dr. William L. Hamilton:

Dr. Lucas N. Amenga-Etego:

### Contents


Supplementary Methods ... 3
Human β-globin genotyping ... 3
PCR reaction mix: ... 3
PCR thermocycler conditions: ... 4
Interpretation of chromatogram traces ... 4
*P. falciparum* genome variant filtering for study population ... 6
Population structure ... 7
Genome Wide Association (GWA) analysis ... 8
Ratios of relative risk of malaria in HbS-carriers based on *Pfsa* genotypes ... 9
FIKK4.2 repeat analysis ... 12
Haplotype analysis at the *Pfsa* loci ... 13
Investigation of recent positive selection at the *Pfsa* loci ... 14
Assessment of power in the analysis of parasite genetic associations with HbC ... 16
Literature cited in supplementary materials ... 19
Supplementary Figures ... 20

### Supplementary Methods

#### Human β-globin genotyping

Human β-globin genotyping at codon 6 followed a PCR amplification and capillary sequencing strategy. The PCR primer sequences used to amplify the locus within the β-globin gene were: acgttggatgGTCTCCTTAAACCTGTCTTG and acgttggatgTCAAACAGACACCATGGTGC. These primers were originally designed for an assay on the Agena/Sequenom MASS array platform [1], which required the 5’ 10bp tails (lower case), though these tails played no role in the current study. The resulting amplicon is 132bp in size. Samples were PCR-amplified and capillary sequenced in 96-well plates. A 3-step PCR approach with graduated annealing temperatures was used (high sequence paralogy at the β-globin locus complicates primer design). PCR amplification used the following reaction mixtures and theromycycling steps:

##### PCR reaction mix:

| Reagent | Quantity (ul) 1x reaction |
|---|---|
| Nuclease Free Water | 10.8 |
| 10x $NH_4$ BioTaq buffer | 2 |
| dNTP mix (10mM each) | 2 |
| $MgCl_2$ (50mM) | 1 |
| Forwards primer (10uM) | 1 |
| Reverse primer (10uM) | 1 |
| Bio Taq polymerase | 0.2 |
| Sample | 2* |
| **Total** | **20** |

Bioline BIOTAQ™ DNA Polymerase (Cat. No. BIO-21060) was used. dNTP mix was made up from Thermo Fisher R0181. The dATP, dGTP, dCTP and dTTP at 100mM was combined and diluted to 10mM each. Primers were ordered from Integrated DNA Technologies (IDT) in 100uM solution and diluted to a working aliquot of 10uM.

*The volume of extracted DNA eluate used for the PCR reactions depended on the sample DNA concentration and volume remaining after parasite sequencing, with nuclease free water (NFW) adjusted accordingly to a final volume of 20ul per 1x reaction. A batch approach was used based on median concentration and volume for each plate. In 4x 96-well plates (356 samples), with a median DNA concentration of 60ng/ul, samples were diluted 1:10 and 2ul of the 10x dilution was used for PCR. For 16x plates (1,483 samples) with a median DNA concentration of 1.7ng/ul, 2ul of sample was used directly for PCR. For 3x plates (297 samples) in which zero (or close to zero) sample volume remained after parasite sequencing (with median DNA concentration 1.3ng/ul), 5ul NFW was added to the wells and 4ul of this was used for PCR. For 1x plate (22 samples), DNA concentration was high (median 62ng/ul) but zero (or close to zero) volume remained post parasite sequencing, so 5ul NFW was added to each well and 2ul of this was used for PCR. 1x plate containing 16 samples could not be located after parasite sequencing so β-globin genotyping was not attempted.

#### PCR thermocycler conditions:

| Reaction step | Description | Temperature (degrees C) | Time |
|---|---|---|---|
| 1 | Initial denaturation | 96 | 1 min |
| 2 | Denaturation | 94 | 45 sec |
| 3 | Annealing | 56 | 45 sec |
| 4 | Extension | 72 | 30 sec |
| Repeat steps 2-4 5x total | | | |
| 5 | Denaturation | 94 | 45 sec |
| 6 | Annealing | 65 | 45 sec |
| 7 | Extension | 72 | 30 sec |
| Repeat steps 5-7 29x total | | | |
| 8 | Final extension | 72 | 10 min |
| 9 | - | 15 | Forever |

Each 96-well plate included a positive and negative control. The positive controls were samples with a known human genotype ($\beta^{A/A}$, $\beta^{A/S}$, $\beta^{S/S}$ or $\beta^{A/C}$), which were anonymous samples selected from the MalariaGEN severe malaria study [2]. The negative control used NFW for the ‘template’. Test wells (between 2 to 4) plus the positive and negative controls were selected from each 96-well plate after PCR and run on 2% agarose gels to check the PCR had been successful. Post-PCR clean-up and capillary sequencing using the reverse primer (acgttggatgTCAAACAGACACCATGGTGC) was performed by Eurofins using the Supreme Run service.

#### Interpretation of chromatogram traces

Capillary sequence chromatogram traces were manually inspected using the 4Peaks software tool. Illustrative screenshots are shown below. All screenshots show the 12 nucleotide sequence CTTCTC**CTC**AGG (as sequenced using the reverse primer, above) – this is the reverse complement of the β-globin sequence CCT**GAG**GAGAAG, where the bold red GAG is the sixth codon, encoding Glutamic acid in the wild-type state. In the images below, variant positions that encode the HbS and HbC haemoglobin proteins are indicated by bold font and underlining.

CTTCTCCTCAGG = $\beta^{A/A}$ genotype, expected to produce HbAA haemoglobin (wild-type homozygote):

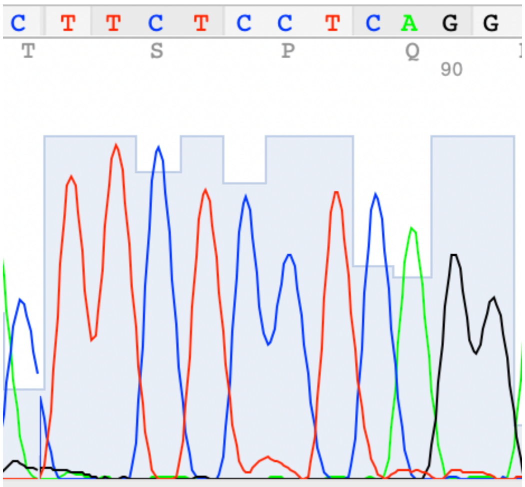


CTTCTCCT(**C/T**)AGG = $\beta^{A/C}$ genotype, expected to produce HbAC haemoglobin (HbC heterozygote):

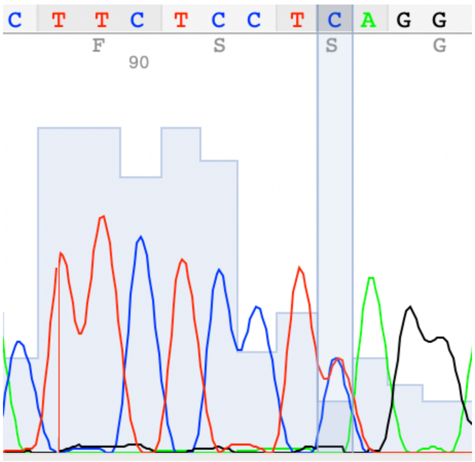


CTTCTCC(**T/A)**CAGG = $\beta^{A/S}$ genotype, expected to produce HbAS haemoglobin (sickle heterozygote):

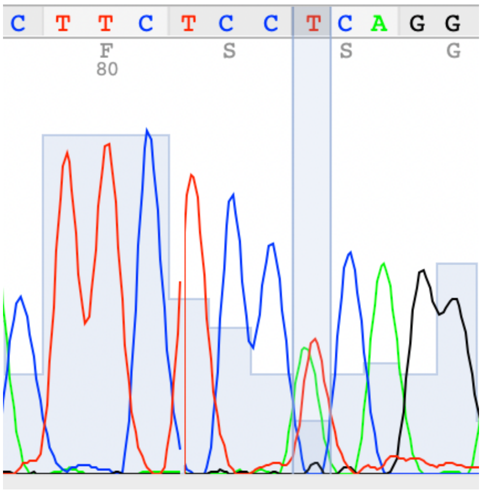


CTTCTCC**A**CAGG = $\beta^{S/S}$ genotype, expected to produce HbSS haemoglobin (sickle homozygote):

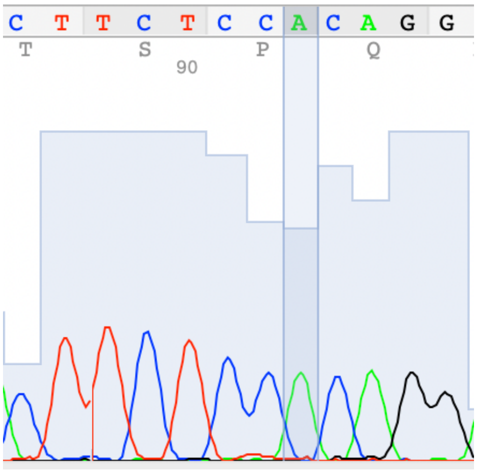


#### *P. falciparum* genome variant filtering for study population

Parasite genomic analyses were conducted using the data structures and curated sample metadata of the MalariaGEN Pf7 data resource [3]. All manipulations of the Pf7 dataset were performed using the Wellcome Sanger Institute High Performance Cluster (HPC). Pf7 includes 10,145,661 variants from 20,864 samples prior to filtering. The following initial filtering steps were applied to streamline the number of variants and reduce the computational requirements for subsequent analysis: biallelic single nucleotide polymorphisms (SNPs) that passed Pf7 quality control (QC) filtering, yielding 2,513,888 SNPs. We explicitly included the previously reported lead associations in the *Pfsa1* and *Pfsa3* regions, which were excluded by the above filters due to their representation in the Pf7 VCF. The Pf7 pipeline calls genotypes as if they were diploid; heterozygous calls represent mixed genotypes and we excluded these from our downstream analysis by treating these genotypes as missing.

To form a working set of variants, we restricted to biallelic SNPs passing Pf7 QC with a minor allele frequency cut-off of 0.1% among the 1,555 samples that passed Pf7 QC, yielding 123,845 SNPs.

#### Population structure

We used principal components analysis (PCA), as implemented in the QCTOOL package [5] to assess population structure in the 1,368 samples passing QC and with both parasite and β-globin genotypes in our data. We restricted attention to the set of 14,352 SNPs with at least 1% minor allele frequency and no more than 25% missing genotypes in these samples, from the 'working set' of 123,845 SNPs described above. PCA works by first estimating a matrix of pairwise relationship values (the "kinship matrix") between all pairs of samples, where $r_{ij}$ denotes the relatedness between sample *i* and *j*.

$$R = (r_{ij})$$

This is scaled so that values close to zero represent little relatedness while values close to 1 indicate complete or near-complete allele sharing, although the precise value of r depends on the sample frequency of the shared alleles. Initial principal components (PCs) revealed a small number of sample pairs with high relatedness values (estimated $r > 0.5$) that dominated the first two principal components (**Supplementary Figure 1A**). We therefore implemented a 'greedy algorithm' to remove one of each sample pair having $r_{ij} > 0.5$. In total we removed 55 samples, leaving 1,313 that were analysed in the logistic regression analysis. We then re-computed principal components and loadings (**Supplementary Figure 1B).** Some population structure was evidenced, reflected in observed variation in the first two principal components which had loadings from SNPs across the genome (**Supplementary Figure 1C**). For comparison, we also computed results using a more stringent relatedness threshold (excluding one of each pair of samples with $r_{ij} > 0.2$); this led to a slightly smaller set of 1,257 samples but did not alter the overall results significantly. To assess the extent to which other measured sample covariates corresponded to PCs, we also plotted the first two principal components coloured by study site, sample year, and HBB genotype (**Supplementary Figure 2**). There was no evidence of population structure being driven by these covariates. Age and sex were not available so were not included as covariates. For the GWA, the first two PCs were included as covariates. For comparison we also computed association test results after controlling for the first 5 PCs, and/or after repeating the analysis using the strict relatedness set of 1,257 samples; there were no significant differences in the results obtained.

In this analysis, principal components 3 and above were observed to have strong loadings from specific loci, including a cluster of ~20 SNPs in the region 38,601 - 39,177 near a Stevor pseudogene (PF3D7_0500600) in PC 3, and a second cluster of ~15 SNPs in the region 1,318,199-1,318,853 near PF3D7_0532600, both on chromosome 5. To reduce the influence of these local regions we repeated the above analysis using a subset of 11,880 SNPs that were thinned for proximity (by iteratively picking a SNP from the 14,352 at random, and then excluding all other SNPs within 100bp either side) and used this version of the PCs for our main analysis.

For the logistic regression GWAS, we analysed variants with a minor allele count of at least 10 among the 1,313 samples that both passed Pf7 QC and had a β-globin genotype call, and after removing 55 samples that drove population structure by PCA (described above). This yielded 18,862 SNPs analysed across 1,313 samples, used for the logistic regression described in the main text.

All genomic data manipulations were performed in Python; genotype and haplotype manipulations and downstream population genetic analyses were performed using the *scikit-allel* package.

We checked whether multiplicity of infection (MOI) differed between infections from individuals carrying different β-globin genotypes, as reflected by Fws [4]. Fws data were extracted for each sample from the publicly available Pf7 MalariaGEN data resource [3]. For HbAA vs HbS-carrying individuals, median Fws was 0.92 (interquartile range, IQR 0.72-1.0) vs. 0.90 (IQR 0.83-1.0), respectively, *P*=0.3448 (Wilcox rank sum test). Therefore, in our dataset we did not find evidence that HbS-carrying individuals tended to have different MOI, on average, compared to HbAA individuals.

#### Genome Wide Association (GWA) analysis

The details of HPTEST have been described previously [6]. Briefly, HPTEST fits a logistic regression model of the parasite genotype (coded as 0 or 1) on the host genotype. For this analysis, we encoded the host genotype also as 0 or 1 (i.e. a dominance model, in which individuals either carry HbS or do not carry HbS). For the HbC main analysis, we used the same approach (in which individuals either carry HbC or do not carry HbC), but we excluded any individuals carrying a sickle allele (HbAS, HbSS, HbSC), to minimise the risk of bias introduced by sickle effects. i.e. we tested for associations between $\beta^{C/no\text{-}S}$ vs $\beta^{A/A}$ individuals. Sample counts for the HbC GWA were: $\beta^{C/no\text{-}S}$, N=315 (303 $\beta^{A/C}$ and 12 $\beta^{C/C}$); vs $\beta^{A/A}$, N=964.

We used the default priors specified in HPTEST, which are Log-F(2,2) – similar to a Gaussian distribution with a variance of 3.5, regarded as a weakly informative prior – for the main genetic effect, and a spread out Log-F – similar to a Gaussian distribution with a variance of 1,600 – for the covariate principal components. Parasite variants in which the minor allele count is less than 10 were excluded. Principal components 1-2 from the PCA were included as covariates to account for population structure. HPTEST computes summaries of the associations for each host and parasite variant including a Bayes Factor (BF, shown in **Figure 2**), an approximate *P*-value, and the regression coefficients and standard deviations. Further information on HPTEST can be found at https://www.well.ox.ac.uk/~gav/hptest/. Frequencies of *Pfsa*+ alleles in sickle and non-sickle individuals are shown in **Supplementary Figure 3**.

#### Ratios of relative risk of malaria in HbS-carriers based on *Pfsa* genotypes

The protective effect of HbS on disease is typically determined by comparing frequencies of HbS in malaria cases and non-malaria population controls. This is commonly quantified as a relative risk (RR). RR is the ratio of the probability of developing malaria in individuals carrying the HbS allele to the probability in those without:

$$RR = \frac{P(disease\ |E = 1)}{P(disease\ |E = 0)}$$

Here, $E = 1$ denotes individuals with HbAS or SS genotypes, and $E = 0$ denotes individuals with the HbAA genotype. The RR is therefore measured relative to a baseline of disease in non-HbS carriers ($E = 0$). An RR less than 1 indicates that individuals with the HbS allele are less likely to develop malaria compared to those without it.

An estimation of how the infecting parasite genotype alters this protection can be determined by comparing HbS frequency to population controls, stratified by possible parasite genotype combinations. The RR of disease observed with a particular parasite genotype $(g)$ in population is:

$$RR_{E=1}(g) = \frac{P\left(disease_g\ |E = 1, S\right)}{P\left(disease_g\ |E = 0, S\right)}$$

Using Bayes Theorem, this can be rearranged into an odds ratio:

$$RR_{E=1}(g) = \frac{P(E = 1|disease_g)}{P(E = 0|disease_g)} \div \frac{P(E = 1|S)}{P(E = 0|S)} = OR$$

This expression implies that $RR_{E=1}(g)$ can be estimated by comparing the frequency of HbS genotypes in a sample of cases involving parasites carrying genotype (first term) to the frequency in a control sample from the general population (second term). An analysis of this type was published with the discovery of the Pfsa loci.

However, in this study no uninfected population controls were collected and so population frequencies are not available for comparison. Therefore, we opted to perform a similar analysis where HbS frequency within cases carrying specific parasite *Pfsa+* genotypes $(g+)$ are instead compared to a baseline of cases caused by Pfsa- infections $(g-)$. Rather than a RR, this calculation produces a "ratio of relative risks" (RRR) between infections of differing parasite genotypes:

$$RRR = \frac{RR_{E=1}(g+)}{RR_{E=1}(g-)}$$

An RRR equal to 1 suggests HbS has similar protective effects across genotypes, while RRR > 1 for a particular genotype suggests that HbS confers less protection against infections with that genotype.

In practice, can then be estimated across parasite genotypes using multinomial logistic regression. To implement this, custom Python code was developed to perform multinomial logistic regression, using as the predictor variable and parasite genotypes at specified *Pfsa* loci as the outcome levels. It is important to note that, because malaria cases in individuals carrying HbS are rare, estimating RRR for certain parasite genotypes using this framework is hindered by small or non-existent sample sizes. To estimate these values for rare parasite genotypes with minimal data and to prevent overfitting of the multinomial model, a Bayesian regularisation method was employed (as described in [6]). This was implemented using Stan with a Gaussian prior on the effect, with a mean of 0 and a standard deviation of 2, and the correlation between parameters across the model set to 0.5. For genotype combinations with low sample counts, the resulting parameter values can be interpreted as plausible estimates given the current data, but larger samples would be required to fully specify these effects.

Estimated effect size for the association between HbS and Pfsa loci, controlling for other Pfsa loci, is shown in **Supplementary Figure 4**. Relative risk of infection in HbS carriers within infections carrying *Pfsa4+* mutations alone (without *Pfsa1+* and *Pfsa3+* alleles) is roughly 3.5-fold greater than in *Pfsa-* infections (**Supplementary Figure 5**). This holds true for non-regularised estimates, suggesting that this result is not an artefact of the implemented regularisation method. Furthermore, the RRR is also increased within infections carrying

*Pfsa1+, Pfsa3*+ and *Pfsa4*+ alleles compared to those carrying only *Pfsa1*+ and *Pfsa3*+ (**Supplementary Figure 5**). The figures plotted in **Supplementary Figure 5** are shown below:

| | | | | ***Regularised estimates*** | | | ***Non-regularised estimates*** | | |
|---|---|---|---|---|---|---|---|---|---|
| ***level*** | **Sample size** | **Number HbS** | **Pfsa genotype** | **OR** | **Lower bound** | **Upper bound** | **OR** | **Lower bound** | **Upper bound** |
| *1* | 1121 | 15 | (0, 0, 0) | | | | | | |
| *2* | 19 | 1 | (0, 0, 1) | 3.56 | 0.77 | 11.27 | 4.10 | 0.51 | 32.70 |
| *3* | 9 | 1 | (0, 1, 0) | 2.87 | 0.48 | 22.65 | 9.22 | 1.08 | 78.37 |
| *4* | 5 | 1 | (0, 1, 1) | 4.47 | 1.72 | 12.41 | 18.43 | 1.94 | 174.85 |
| *5* | 3 | 0 | (1, 0, 0) | 2.28 | 0.13 | 23.53 | | | |
| *6* | 2 | 0 | (1, 0, 1) | 6.32 | 0.85 | 30.43 | | | |
| *7* | 17 | 6 | (1, 1, 0) | 23.87 | 9.85 | 100.81 | 40.22 | 13.15 | 122.97 |
| *8* | 19 | 7 | (1, 1, 1) | 29.86 | 12.73 | 87.01 | 43.01 | 14.87 | 124.43 |

In the table above, a zero indicates cases carry a *Pfsa*- allele at the locus and 1 indicates a *Pfsa*+ allele. Estimates of odds ratio (OR) and 95% confidence intervals made with and without using Bayesian regularisation are given.

Taken together this suggests an independent effect of the *Pfsa4*+ allele in improving the ability of *P. falciparum* parasites to infect HbS individuals. This analysis revealed differences in RRR between the present study in Ghana and previous studies in Kenya and Gambia [6]. In Ghana, the RRR for infections with *Pfsa1, Pfsa3,* and *Pfsa4*+ alleles was 30, indicating that HbS carriers have a 30-fold higher relative risk when infected with *Pfsa*+ parasites compared to *Pfsa*- infections. This contrasts with findings from Kenya and Gambia, where the RRR was approximately 64, with HbS carriers infected by *Pfsa1-3*+ genotypes having a 64-times greater risk compared to *Pfsa*- infections (RR≈1 for *Pfsa*+ infections versus RR<1/64th for *Pfsa*- infections, therefore RRR is approximately 64).

These calculations were made relative to matched population controls, which may influence the results. However, they still suggest a varying impact of parasite genetics on HbS-related resistance across populations. Notably, in the Ghanaian cohort, 15 HbS-carrying individuals were infected by *Pfsa*- parasites. This contrasts with the discovery cohorts, where only four such infections were reported in Kenya, and none in Gambia [6]. It is important to again note that these estimates are based on a limited number of malaria cases in HbS carriers.

Additional datasets are required to determine whether these observations reflect true population-level variations or if the estimates will converge with larger sample sizes and improved statistical power.

#### FIKK4.2 repeat analysis

Previous work has identified copy number variation within FIKK4.2 [7] within a 6-residue amino acid motif (hexapeptides) with the majority of repeats being of the form SD[HNS]NH[KM]. We used 16 available *P. falciparum* genome assemblies to investigate this repeat [8], as follows. First, we extracted a 101bp kmer centred at the lead *Pfsa4* SNP (chr4:1,121,472 from the Pf3D7 reference assembly), sequence shown below:

K=ACGATAATGATGACAGTGATGCAAGCGATGCAGTTCATGAAGATATTGAGTTACTTG
AGTCTTATAGTGATTTGAATAAATTTAATGAGATGTTAACAGAA

We used BLAT to align this to the available assemblies. BLAT is a sensitive aligner and we focussed on alignments with at least 80 matching bases. Notably, this kmer aligned to chromosome 4 with >= 99 matching bases on all assemblies, except for the PfSD01 isolate from Sudan, for which it aligned to chromosome 7 with 99 matching bases (but not to chromosome 4). This is consistent with the publicly available gene annotations for this genome, and suggests this locus has undergone translocation in some isolates.

We then expanded the alignment coordinates by adding 10kb on either side and extracted DNA sequence from each assembly, and computed a multiple sequence alignment (MSA) using MAFFTv7.490. Using this MSA, we identified a set of DNA kmers which are shared identically across all isolates and mark the start and end of the coding sequence of FIKK4.2 in each exon (detailed in the table below).

| Exon | 5' kmer (DNA/*AA sequence*) | 3' kmer |
|---|---|---|
| 1 | ATGAATTATTTTTCTAAATACAAAGTTATT<br>*M N Y F S K Y K V I* | TATTTTTTGTTTATAATTCCATTG<br>*Y F L F I I P L* |
| 2 | AATGAAGTAATATACAATAAATAT<br>*N Q V I Y N K Y* | GGTCTCAAAGATATAATTAAC<br>*G L K D I I N* |
| 3 | AAATTATTAAGCCTAGAAAGT<br>*K L L S L Q S* | GAACATCCATGGTGGATTAATGAAGATTAA<br>Q H P W W I N Q D * |

We then extracted the full amino acid sequence for FIKK4.2 from each assembly, by using the above kmers to extract the coding sequence. To investigate the repeat region, we plotted DNA kmer (k=31) sharing between 3D7 and all other assemblies (**Supplementary Figure 6**). In 3D7, the repeat unit ends at position 1,121,413, which is only 59 bases away from the lead *Pfsa4* SNP. To evaluate the length of the hexamer repeat, we also used the multiple sequence alignment to identify short amino acid sequences shared between assemblies and flanking the hexamer repeats. This appears of the form EEDKNM[6 residue repeats]NNNNKD. We used this pattern to extract the repeat sequence from each amino acid sequence and to count repeat units.

#### Haplotype analysis at the *Pfsa* loci

In principle it is possible that the *Pfsa*+ mutations arose independently in separate locations, or that they may have a shared evolutionary origin and have spread between populations. To assess this, we created images of haplotypes at the *Pfsa1-4* regions across populations in Africa.

For this analysis we opted to use the entire MalariaGEN Pf7 resource which was sampled from global populations [3]. To create a curated African-specific set of genotypes for this analysis, VCF files (representing genetic variant sites and genotypes) were downloaded from the Pf7 resource page (https://www.malariagen.net/resource/34/) and post-processed using the same method as described in [9]. Briefly, samples were restricted to those from African study sites, Pf7 QC pass, and highly clonal (Fws>0.9) (N=4788 samples). Variants were filtered to Pf7 QC pass. We use the vcflib 'vcfilter' tool to remove alleles with fewer than 20 copies at multi-allelic sites (across the 4,788 samples), effectively converting many such variants to bi-allelic, and we retained all remaining bi-allelic sites yielding 2,959,655 variants with reference and alternate allele assignments. Heterozygous calls were replaced with missing genotypes and then re-imputed in haploid form using beagle (version 5.4, beagle.01Mar24.d36.jar), with a small number of remaining heterozygous calls (5486) replaced with reference homozygous genotypes. Finally, QCTOOL was used to "polarise" alleles in the VCF to represent ancestral and derived states (i.e. so that the ancestral allele is the first allele, and the derived allele is the second allele). The determination of ancestral and derived alleles follows the same methodology as described in [9].

To analyse patterns of haplotype sharing at the *Pfsa* loci, we focussed on a 10kb centred on the lead sickle-associated variant (identified as the variant with the highest Bayes factor) at each of the *Pfsa1-3* loci. At *Pfsa4*, we adjusted this to only include 7kb due to the higher density of variants at this locus.

Genotypes were parsed into python code using the package *scikit-allel* (version 1.3.11) and clustered using the package *scipy* (version 1.14.1). Initially, the haplotype matrix was transposed so that each row represented a sample, and each column represented a variant. Hierarchical clustering was then performed using the average linkage method, which calculated the distance between clusters as the average distance between all pairs of genotypes in the clusters. The function used the resulting dendrogram to determine the order of the samples, reordered the haplotype matrix and the samples array accordingly, and returned the reordered haplotype matrix and samples array.

Haplotypes carrying the *Pfsa*+ allele at the *Pfsa1, Pfsa2, Pfsa3* and *Pfsa4* loci appeared distinct from those carrying the *Pfsa-* allele (**Supplementary Figure 7**), causing haplotypes to be clustered by Pfsa+ genotype rather than by population. Furthermore, at all the loci several other derived mutations appeared to be essentially perfectly correlated to the *Pfsa*+ site (e.g. at *Pfsa1* Pf3D7_02_v3:630150 and Pf3D7_02_v3:630290, at *Pfsa2* Pf3D7_02_v3:813895-814329, at *Pfsa3*, Pf3D7_11_v3:1054282 and Pf3D7_11_v3:1054587 and at *Pfsa4* Pf3D7_04_v3:1121457 and 1119762).

At each locus there were also derived mutations which appeared on a subset of *Pfsa*+ haplotypes (but were still not seen on *Pfsa* haplotypes). For example, at the *Pfsa1* region some *Pfsa1*+ parasites carried the derived allele at the Pf3D7 02 v3:630420 locus whereas others did not. These features strongly indicate that haplotypes at each locus have a single mutational origin as it is highly unlikely that further mutations would coincide with the central *Pfsa*+ allele independently within multiple evolutionary lineages. Following the *Pfsa*+ mutation, it appears that other mutations have occurred within these lineages leading to modern *Pfsa*+ haplotypes. Taken together, this suggests that these haplotypes are shared across Africa and thus have a potentially ancient evolutionary origin. It should be noted that structural variation has been previously noted at the *Pfsa3* locus [6]; structural variation has not been taken account of in this analysis, and therefore interpretation of genealogy at this locus may be hindered.

#### Investigation of recent positive selection at the *Pfsa* loci

To assess for extended haplotypes surrounding the *Pfsa* loci that may indicate recent positive selection [10, 11], we used a common metric of haplotype sharing, extended haplotype homozygosity (EHH). Starting at a focal allele A, EHH at a position x on the chromosome is defined as the probability that two randomly chosen haplotypes both carrying A are identical by descent (IBD) across the region between A and x. EHH starts at 1 near the focal allele, and decays as x moves away from a focal locus, due to recombination. The area under the EHH curve is known as the 'integrated haplotype homozygosity' (iHH). In practice, iHH for the ancestral allele is usually compared to iHH for the derived allele at any locus, controlling for local variation in recombination or mutation rates, leading to the 'un-normalised IHS' metric:

$$uIHS = log\frac{iHH_{derived}}{iHH_{ancestral}}$$

Under pure genetic drift, the value of uiHS will depend on the allele frequency (since, typically, rarer alleles tend to be younger and therefore have larger iHH values). Therefore, to detect selected alleles it is usual to work with uiHS values within allele frequency bins (iHS values) [12].

For simplicity in our analysis, the filtered variant call set developed from the Pf7 resource which contained entirely homozygous non-missing haplotype calls was used, which had been polarised to represent derived and ancestral alleles [9]. In our implementation, EHH, iHH and iHS was calculated across the *P. falciparum* genome over all biallelic SNPs with a minimum minor allele frequency of 5%. Genomes were re-formatted into '.hap' and '.map' file formats using custom code and EHH and iHS was then calculated for every variant across each chromosome using the R package rehh within 2.5% allele frequency bins separately for each population [13]. Populations were limited to those in Africa which contained at least 50 samples (Ghana (N=1582), Gambia (N=574), Cameroon (N=143), Mali (N=691), Sudan (N=65), Guinea (N=84), Kenya (N=401), Mauritania (N=58), Nigeria (N=83), DRC (N=259), Benin (N=101), Malawi (N=114), Senegal (N=118) and Tanzania (N=345)). Data was treated as 'phased' by rehh (as each *P. falciparum* whole genome sequence had been simplified to a haploid form).

iHS values within each population at lead *Pfsa1-4*+ mutations and two previously identified chloroquine drug resistance loci in the CRT gene (Pf3D7_07_v3:403625:A>C) [14] and AAT1 gene (Pf3D7_06_v3:1215233:G>A) [15] are available in **supplementary table 4**. At the *Pfsa3* locus, a nearby variant in strong linkage with the previously identified lead mutation was utilized for analysis, as it was present as a biallelic variant within the Pf7 variant call dataset. The lead variant was excluded from the analysis due to the presence of a rare deletion at this locus (Pf3D7_11_v3:1058035) causing the entry in the variant call file to be multiallelic.

In these calculations, large negative iHS values indicate an extended haplotype carrying the derived allele, suggesting the occurrence of positive selection for the derived allele, whereas unusually positive values indicate a short haplotype.

Both drug resistant loci showed evidence of positive selection with strongly negative iHS values and *P*-values < 0.05 within most tested populations. This contrasts to all four *Pfsa* loci which all showed little evidence for positive selection with slightly negative (iHS≈-1) or neutral values (iHS≈0) in every tested population. Overall, these results suggest that the *Pfsa* regions are unlikely to have undergone a recent selective sweep in any of the studied populations.

#### Assessment of power in the analysis of parasite genetic associations with HbC

In the main text we report no associations of parasite genotypes with the HbC resistance allele. There is a possibility that this is due to insufficient power within our study. Although HbC is more common within our sample compared to HbS ($F^{HbC}$= 12.4%, $F^{HbS}$ = 1.4%), the effect of HbS genotype on parasite genotype at the *Pfsa* loci is large (OR > 4) such that effects are detectable. Should the effect of HbC on parasite genotype be more subtle, this could strongly diminish power.

To test this, we conducted a power analysis using simulations. For a population of individuals with matched HbC genotypes to our sample (that is, 1008 carrying the HbAA genotype, 312 carrying HbAC and 12 carrying HbCC), parasite genotypes were simulated under varying effect sizes ($e^{\beta}$=0.25, 0.5, 2 or 4) and underlying parasite population allele frequencies ($F$ = 5%, 10%, 25% or 50%).

Let $g^{AA}$, $g^{AC}$ and $g^{CC}$represent the proportions of HbAA, HbAC and HbCC individuals within our sample. Parasite allele odds within HbAA individuals ($e^{\mu}$), HbAC individuals ($e^{\mu+\beta}$) and HbCC individuals ($e^{\mu+2\beta}$) was computed assuming an additive effect using the following formula:

$$e^{\mu} = \frac{F}{g^{AA} + g^{AC}e^{\beta} + g^{CC}e^{2\beta}}$$

This value was then used to compute odds in HbAC and HbCC individuals for a given $\beta$ and converted to a probability using x → x / (1 + x). Using these values 1000 sets of genotypes were simulated using a binomial distribution.

Using the generated sets of genotype calls, a logistic regression was then performed using human genotype as the predictor and parasite genotype as the outcome. HbAC and HbCC genotypes were grouped together (i.e. a dominant effect) to reflect that used within the study. For a *P*-value threshold denoted $T$ , given that an association is true (represented as $A$ ), the power of a test of association can be expressed as P(P < T | A). Therefore, for a range of *P*-

value thresholds, the proportion of logistic regression association results where p < T represents the power.

The results of this analysis are presented in the figure below.

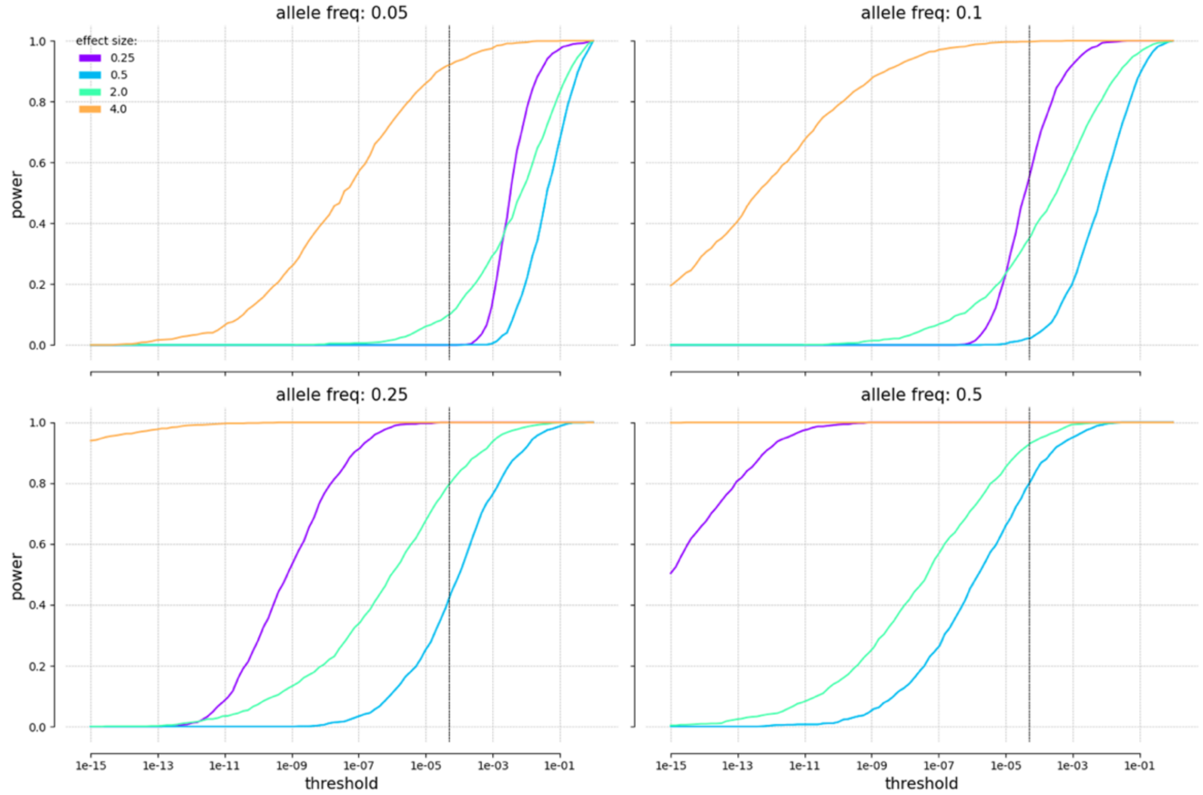


Power, calculated using simulations, is displayed on the y-axis, with different *P*-value thresholds (log10) shown on the x-axis. These values were computed from 1000 sets of parasite genotypes simulated as conditional on human genotypes (HbAA, HbAC, or HbCC), assuming an additive effect of the HbC allele on parasite genotype. Five baseline allele frequencies (representing the overall population frequency of the parasite allele) were tested: 5%, 10%, 25%, and 50%. Four effect size magnitudes (OR = 0.25, 0.5, 2, and 4) are indicated by the colour of the plotted lines. A *P*-value threshold at the Bonferroni correction level ($5 \times 10^{-5}$) is marked with a dashed black line.

Although we do not utilise a *P*-value threshold within this research, a reasonable threshold could be approximately Bonferroni-corrected (1/19726 $\approx$ $5 \times 10^{-5}$) which has been highlighted on the plot as a dashed line. This indicates that we are well-powered to detect strong effects at magnitudes similar to that of HbS-Pfsa (OR ≥ 4) at any allele frequency. However, the power to detect less pronounced frequency-increasing effects (OR = 2) or strong frequency-reducing effects (OR = 0.25) is minimal for rare parasite alleles. The power to detect moderate effects with an OR = 2 at a 10% allele frequency is less than 40%. Power improves for this effect size in more common variants with an allele frequency of 25% or greater.

Therefore, if a parasite genetic association with HbC exists but is more subtle than that of HbS-Pfsa, it is unlikely to be detected unless it is common. This highlights the requirement for further datasets including HbC typing in order to ascertain whether parasite evolution in response to this resistance mutation has occurred.

### Supplementary Figures

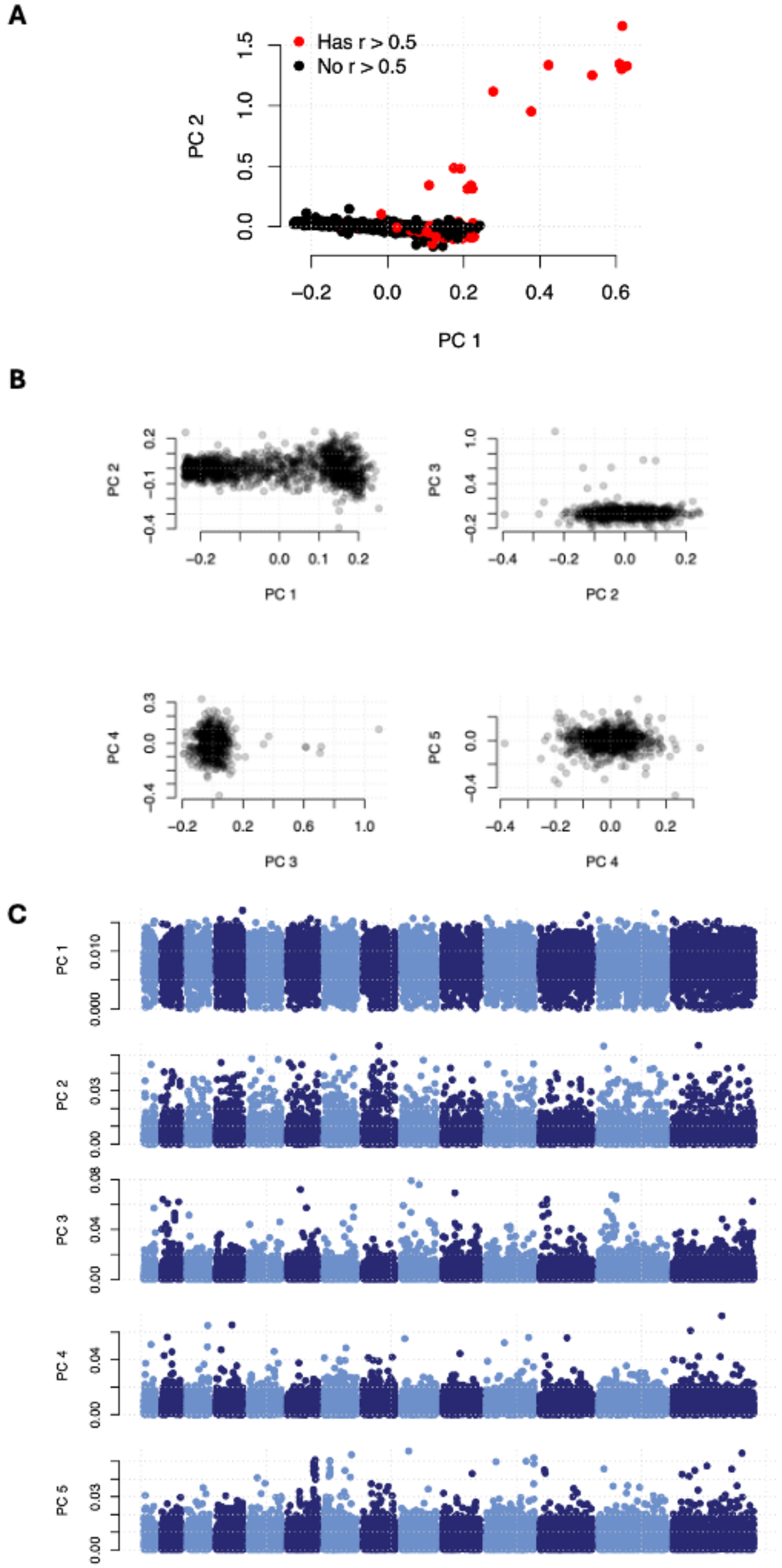

**Supplementary Figure 1**. Assessments for population structure among the 1,368 samples with both a β-globin genotype call and high-quality *P. falciparum* whole genome sequence from the MalariaGEN Pf7 data resource. Principal Components Analysis (PCA) was calculated using genome-wide SNPs as described in supplementary methods. (**A**) Plot shows the first and second principal components from the PCA of all 1,368 available samples, highlighting 55 samples (in red) with high relatedness that were removed to yield the 1,313 samples included in the genome-wide association (GWA) analysis by logistic regression. (**B**) Plots showing principal components 1-5 after re-computing the PCA with the 55 highly related samples removed (N=1,313). (**C**) Genome-wide association for each principal component of the PCA for the 1,313 samples after removing 55 closely related samples.

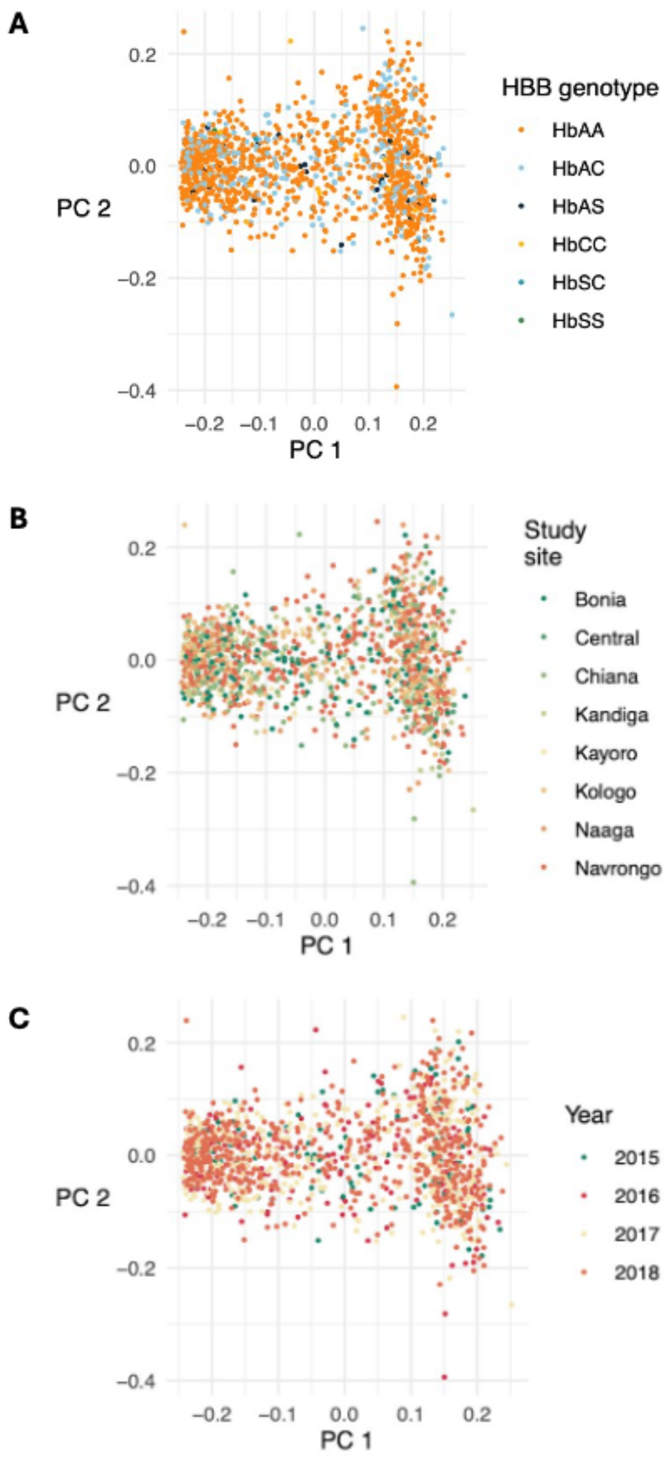


**Supplementary Figure 2**. PCA plots of *P. falciparum* samples included in the study, coloured by GWAS covariates: (A) HBB genotype; (B) Study site; (C) Year of collection. There was no evidence of population structure according to any of these covariates.

A

B

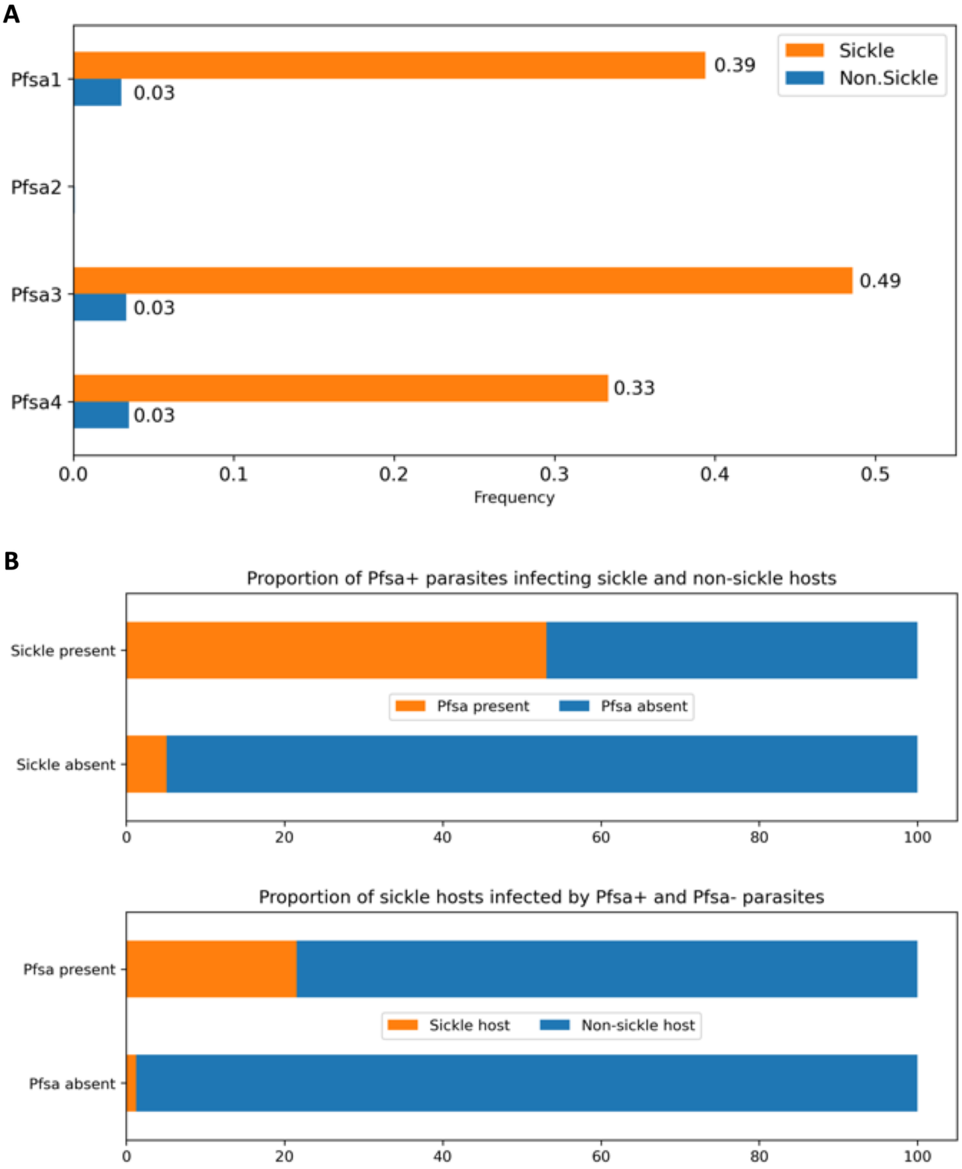


**Supplementary Figure 3**. Frequencies of *Pfsa*+ alleles in sickle and non-sickle individuals. (**A**) Frequencies of each of the four *Pfsa*+ alleles in sickle and non-sickle individuals, for the Ghanaian population with mild malaria reported in this study. *Pfsa2*+ was absent from this population. Note that the *Pfsa4*+ allele is present in the 3D7 reference. (**B**) Proportion of parasites with at least one *Pfsa1-4*+ allele infecting sickle and non-sickle individuals (top), and proportion of people with and without sickle infected by parasites with or without *Pfsa*+ alleles (**bottom**), from this study population. Missing or heterozygous parasite genotypes at the *Pfsa* loci were excluded. To count as '*Pfsa* present', at least one *Pfsa*+ allele had to be present in a sample (*Pfsa1* = chr2:631,190; *Pfsa2* = chr2:814,288; *Pfsa3* = chr11:1,058,035; *Pfsa4* (newly reported in this study) = chr4:1,121,472). Despite the overall prevalence of sickle alleles only being around 3% in this cohort, and only around 4% of parasites carrying *Pfsa*+ alleles, over half of the people carrying sickle alleles were infected by parasites with at least one *Pfsa*+ allele. From the parasite's perspective, over 20% of infections caused by parasites carrying at least one *Pfsa*+ allele occur in individuals carrying a sickle allele.

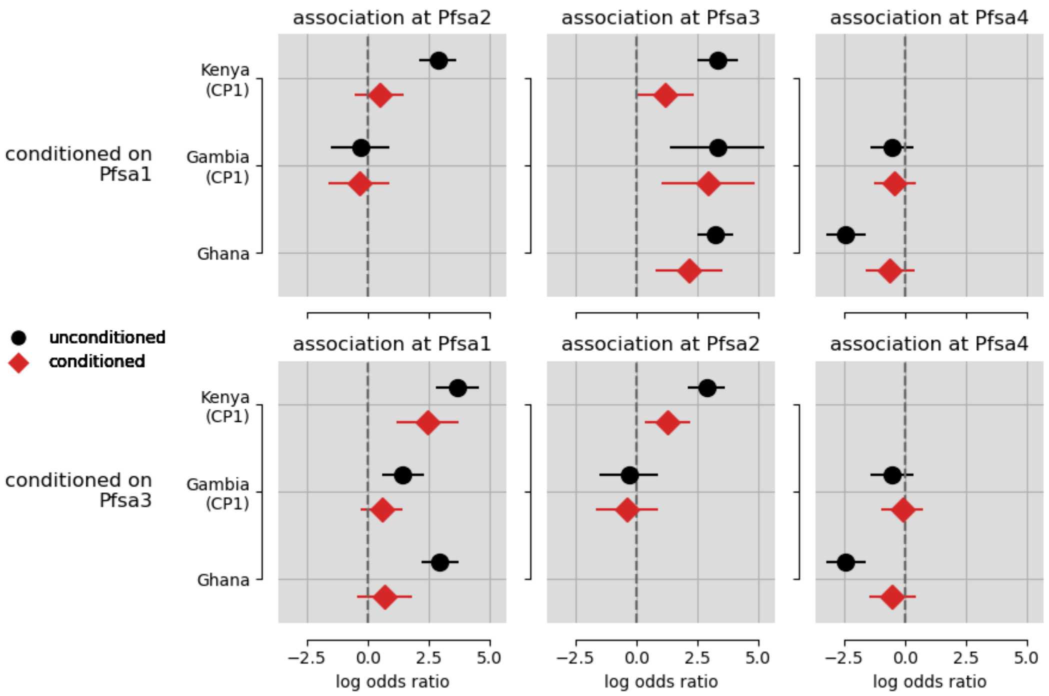


**Supplementary Figure 4.** Estimated effect size (shown as a log odd's ratio and 95% confidence interval) for the association between HbS and *Pfsa* loci with (red diamond) and without (black circle) the inclusion of genotype at the *Pfsa1* (top row) and *Pfsa3* (bottom row) loci as covariates in the association. The locus of the tested association is indicated by the title of each subplot.

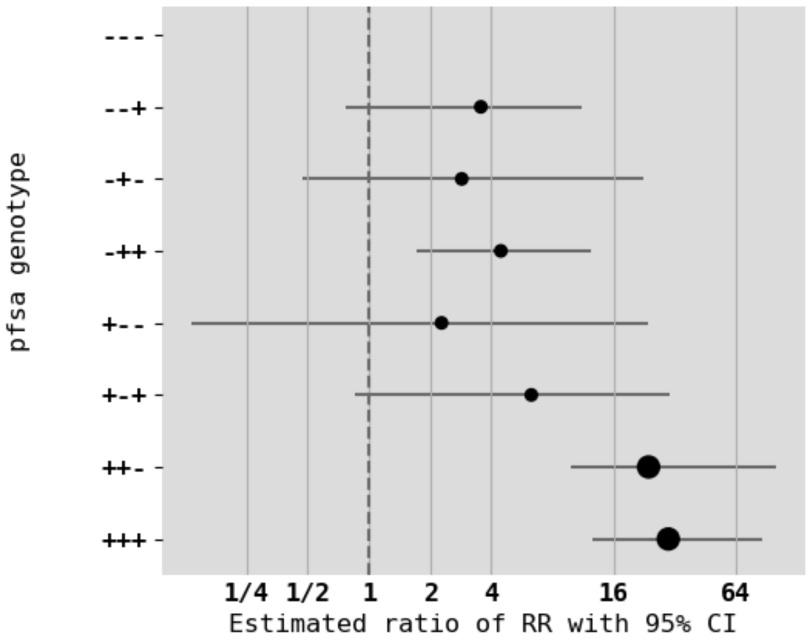


**Supplementary Figure 5.** Estimated relative risk of severe malaria for HbS carriers in infections of different *Pfsa* genotypes. Ratio of relative risk (RRR) is stratified by infecting malaria parasite genotype at the *Pfsa1, Pfsa3* and *Pfsa4* loci. RRRs were calculated using a multinomial logistic regression using infections with *Pfsa-* parasites as a baseline. A '+' for the *Pfsa* genotype indicates possessing the effect allele. Larger points indicate where a sample of greater than five infections in HbS carrying individuals was present. To avoid over-fitting of points with small sample sizes, Bayesian regularisation was performed using a weakly informative Gaussian prior with mean 0 and standard deviation of 2 for each parameter, and between-parameter correlation set to 0.5.

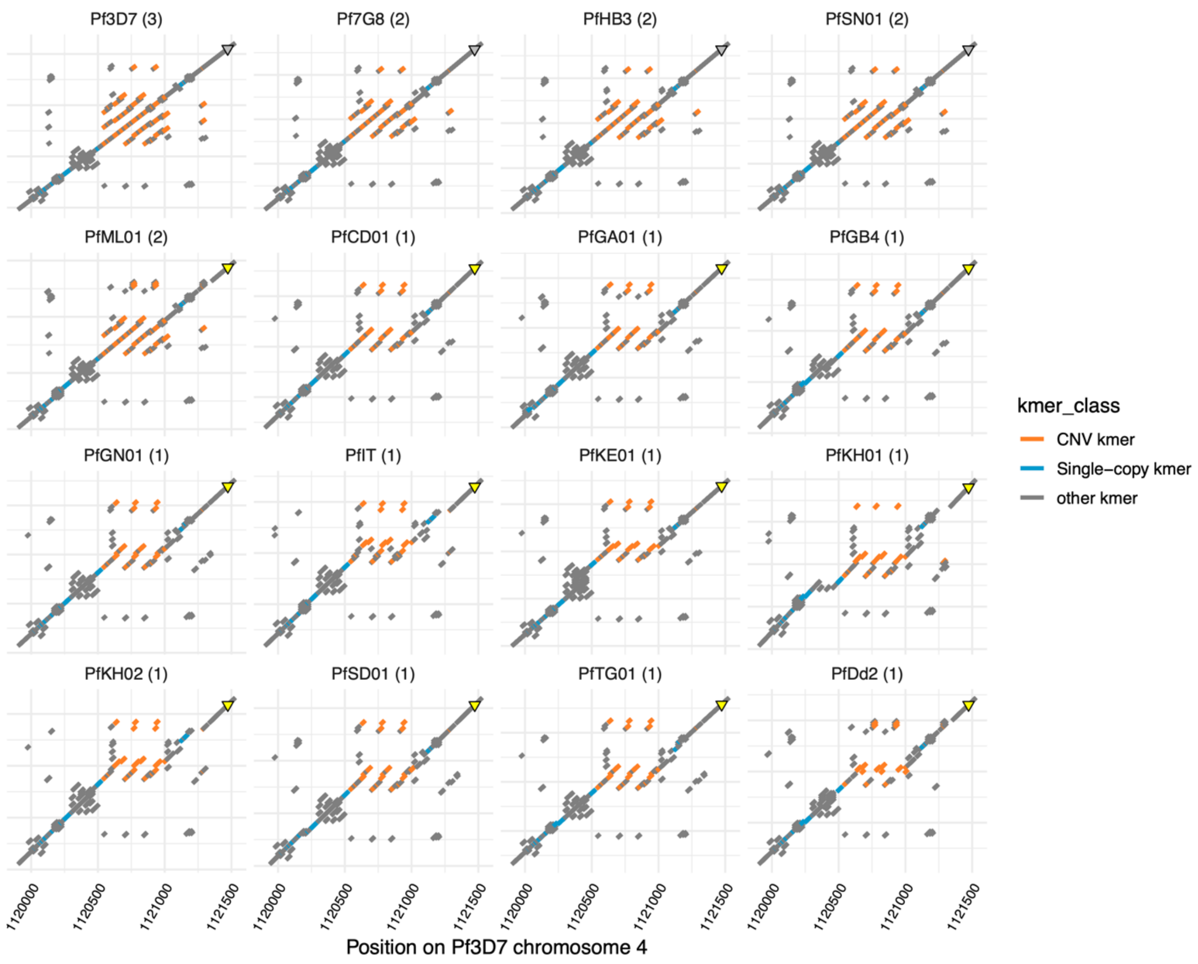


**Supplementary Figure 6**. Detail of repeat structure in available reference assemblies. The figure focusses on a region around the FIKK4.2 hexamer repeat. In each panel, points show 31bp DNA kmer that is shared identically between the Pf3D7 genome (x axis) and the indicated genome (y axis). Orange points indicate kmers in the region 1,120,000-1,121,250 that appear exactly three times in Pf3D7, but less than three times in at least one assembly. Blue points indicate kmers that appear exactly once in each assembly. The triangle indicates the lead *Pfsa4* SNP position; coloured in grey if the *Pfsa4+* (sickle-associated) allele is present, and yellow if the *Pfsa4-* (not sickle-associated) allele is present. The isolate names are shown above each plot with the copy number in brackets.

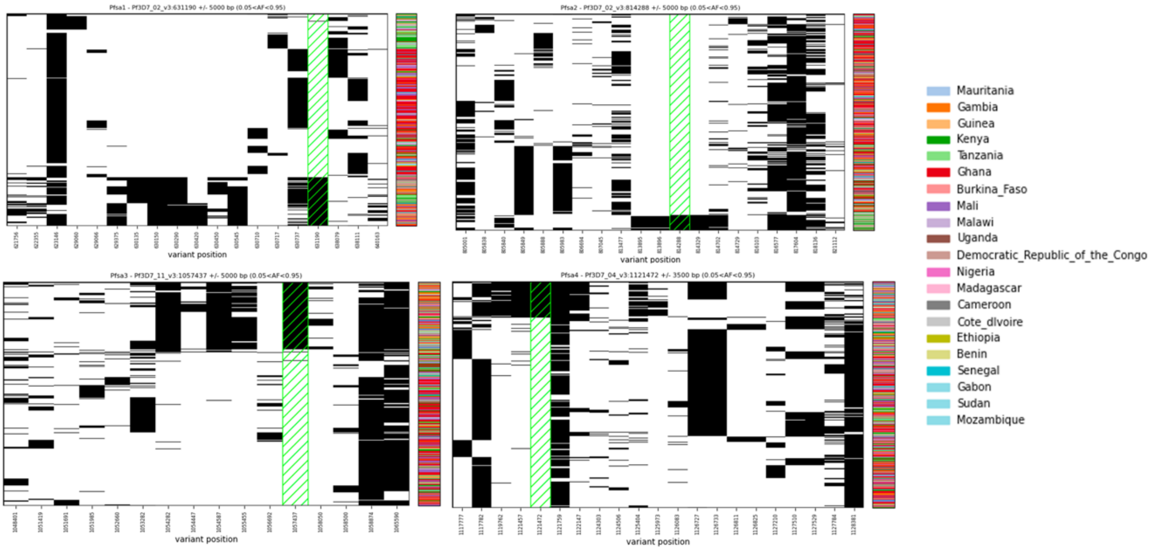


**Supplementary Figure 7**. Haplotypes at the *Pfsa* loci in samples from the MalariaGEN Pf7 resource. Haplotypes at the previously discovered *Pfsa1-4* loci are shown. SNPs with an allele frequency <0.05 or >0.95 were filtered prior to clustering. A colour bar indicating the country of origin of the sample is shown on the right. The top HbS-associated variant at each locus is highlighted in hashed neon green. Haplotypes carrying the *Pfsa+* allele at the *Pfsa1, Pfsa2, Pfsa3* and *Pfsa4* loci appear distinct from those carrying the *Pfsa-* allele, causing haplotypes to be clustered by *Pfsa+* genotype rather than by population, consistent with a single evolutionary origin for each *Pfsa+* allele.

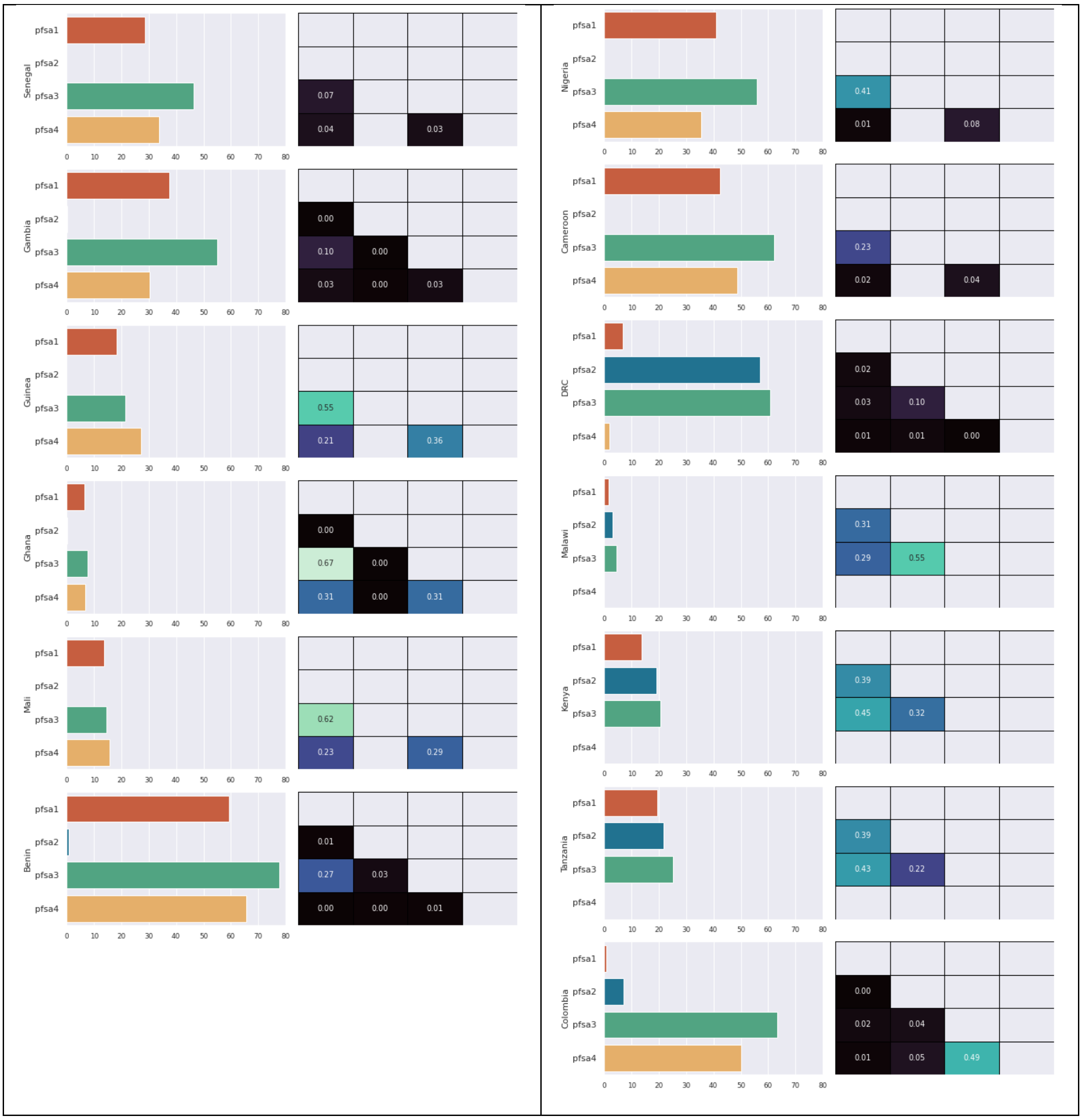


**Supplementary Figure 8**. *Pfsa1-4+* allele frequencies and linkage disequilibrium (LD) for all African and South American countries included in the MalariaGEN Pf7 data resource with ≥100 samples that pass QC. Asian countries were excluded because the frequencies of the *Pfsa+* alleles are extremely low.
